# SHORT-TERM CALORIC RESTRICTION IN MICE PROMOTES RESOLUTION OF ATHEROSCLEROSIS, WHILE WEIGHT REGAIN ACCELERATES ITS PROGRESSION

**DOI:** 10.1101/2023.05.07.539777

**Authors:** Bianca Scolaro, Emily J. Brown, Franziska Krautter, Marie Petitjean, Casey Donahoe, Stephanie Pena, Michela L. Garabedian, Cyrus A. Nikain, Maria Laskou, Ozlem Tufanli, Carmen Hannemann, Myriam Aouadi, Ada Weinstock, Edward A. Fisher

## Abstract

While weight loss is highly recommended for those with obesity, >60% will regain their lost weight. This weight cycling is associated with elevated risk of cardiovascular disease, relative to never having lost weight. How weight loss/regain *directly* influence atherosclerotic inflammation is unknown. Thus, we studied short-term caloric restriction (stCR) in obese hypercholesterolemic mice, without confounding effects from changes in diet composition. Weight loss was found to promote atherosclerosis resolution independent of plasma cholesterol. From single-cell RNA-sequencing and subsequent mechanistic studies, this can be partly attributed to a unique subset of macrophages accumulating with stCR in epididymal adipose tissue (eWAT) and atherosclerotic plaques. These macrophages, distinguished by high expression of *Fcgr4*, help to clear necrotic cores in atherosclerotic plaques. Conversely, weight regain (WR) following stCR accelerated atherosclerosis progression with disappearance of Fcgr4+ macrophages from eWAT and plaques. Furthermore, WR caused reprogramming of immune progenitors, sustaining hyper-inflammatory responsiveness. In summary, we have developed a model to investigate the inflammatory effects of weight cycling on atherosclerosis and the interplay between adipose tissue, bone marrow, and plaques. The findings suggest potential approaches to promote atherosclerotic plaque resolution in obesity and weight cycling through induction of Fcgr4+ macrophages and inhibition of immune progenitor reprogramming.

**Graphical abstract:** 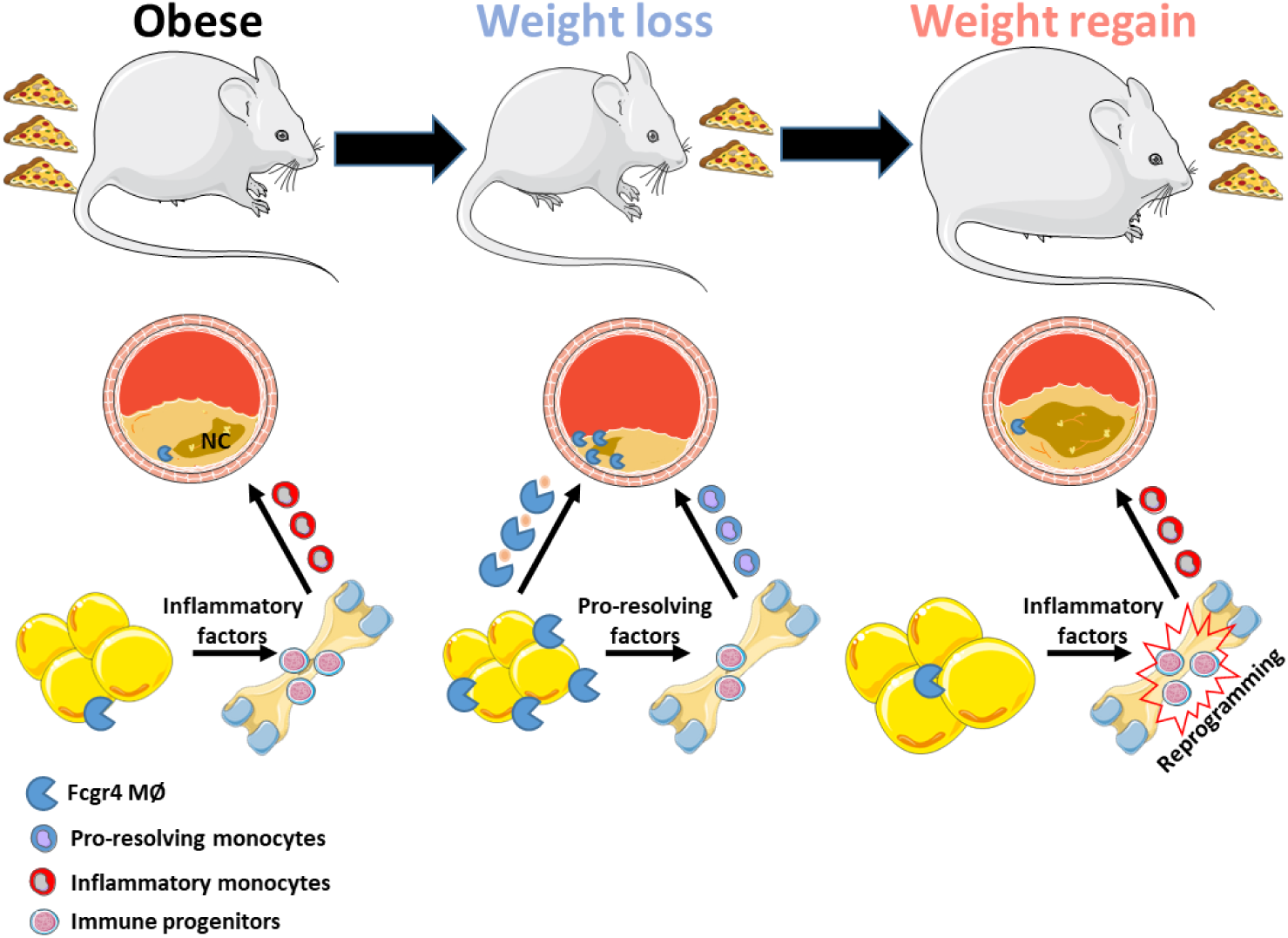

## INTRODUCTION

Obesity contributes to the establishment and progression of many diseases, including the leading cause of death, atherosclerotic cardiovascular disease (CVD(1, 2)). A major factor thought to underlie this association is the heightened production and release of a variety of potent inflammatory factors (presumably secreted from visceral adipose tissue), such as interleukin (IL)-6, IL-1ꞵ, S100A8/9, and TNFα, based on observational studies in humans and mouse models (reviewed in(3)). Contributing to these inflammatory changes are the effects of obesity on insulin resistance and glucose homeostasis. These metabolic perturbations are potent stimulators of local and systemic inflammation through, for example, the production of reactive oxygen species, ligands for the receptor of advanced glycation end-products, and activation of macrophages. In mouse studies, it has also been shown that obesity has effects on hematopoiesis(4). This appears to be through IL-1ꞵ secretion from visceral adipose tissue (VAT) macrophages, which is promoted by neighboring adipocytes. This subsequently stimulates the proliferation of bone marrow precursors of monocytes and neutrophils (the major mediators of innate inflammatory responses), thereby increasing their abundance in the circulation and VAT(4).

Sustained weight loss can decrease the risk or severity of many obesity-associated diseases, including CVD(5–7). In addition to improving insulin sensitivity and glucose homeostasis(8), weight loss decreases the leukocytosis and increases in inflammatory markers associated with obesity(9, 10). Furthermore, weight regain (which occurs in >60% of dieters) increases clinical CVD risk(11) and adipose tissue inflammation in mouse models(12) over what is observed in those never having lost weight. However, cellular and molecular immune mechanisms that facilitate resolution of obesity-related inflammation with CR and heighten inflammation with weight regain are incompletely known.

To begin addressing these gaps, we previously performed(13) single-cell RNA sequencing (scRNAseq) of epididymal white adipose tissue (eWAT; a surrogate in mice for VAT) because of the above noted association between this depot and adverse consequences of obesity on inflammation(14). We profiled eWAT leukocytes of mice fed a high fat high cholesterol (HFHC) diet to induce obesity before and after short term (2 weeks) reduction of caloric intake by 30%(13). By maintaining HFHC feeding throughout the study and reducing caloric intake, we achieved weight loss while avoiding confounding the data by inducing epigenetic changes in monocytes and their precursors related to diet compositional changes that result by shifting to chow(15, 16). Our results demonstrated that this type of CR (which we term stCR, for short-term CR) induced a unique immune milieu in the eWAT, neither totally resembling the obese nor the lean landscape.

Most notable was finding a novel adipose tissue macrophage (ATM) population that accumulates with stCR and was characterized by high expression of the IgG antibody receptor *Fcgr4* (in humans, *Fcgr3a*), which promotes phagocytosis (reviewed in (17)). This ATM population expressed many other genes also implicated in phagocytosis, so we hypothesized that these cells assist in clearing apoptotic cells, products of shrinking adipose tissue, and, indeed, there was evidence of this(13). Clearance of apoptotic cells by macrophages (a process termed efferocytosis) is a critical process in inflammation resolution(18–20), so we hypothesized that stCR may induce this process in other macrophage-rich tissues as well, and thereby promote inflammation resolution.

In human atherosclerotic plaques, the necrotic core, an area where apoptotic and necrotic cells accumulate, expands as disease progresses. This results in plaque vulnerability, which can lead to rupture and myocardial infarction(21). Mechanistic studies in mice have shown that the expansion of the necrotic core is a result of ongoing cell death coupled with inadequate efferocytosis, contributing to a failure to resolve plaque inflammation (reviewed in(22, 23)). Based on studies of mouse models of atherosclerosis regression (e.g., (24–26)), other features of plaque inflammation resolution include an overall decrease of macrophage abundance, higher expression of inflammation pro-resolving genes (e.g. *Mrc1*, *Arg1*, *CD163*) in the remaining plaque macrophages, and increased plaque collagen content(24–26).

Taking into consideration our previous results in eWAT(13) and their potential relevance to atherosclerosis, we aimed at elucidating the effects of stCR on established atherosclerosis, especially on plaque inflammation and the necrotic core. We hypothesized that stCR promotes simultaneous benefits in both eWAT and atherosclerotic plaques by enhancing clearance of dying/dead cells and inflammation resolution(13). We also sought to recapitulate, in mice, findings consistent with the clinical observations of increased weight following weight loss (i.e. weight cycling), and if so, to determine its mechanisms. In both diet settings (stCR-induced weight loss and weight regain), we were particularly interested in the roles of Fcgr4+ macrophages, given their aforementioned association with the beneficial changes observed in the eWAT studies(13). We hypothesized that the abundance of these cells would vary in the metabolic states (obese, stCR, weight regain) and that they would have functional beneficial properties in both eWAT and plaques.

We successfully established mouse models relevant to these important clinical issues in obesity and atherosclerosis with which to test our hypotheses. As will be presented, the application of an array of modes of analyses, including extensive bioinformatics, has revealed several insights into the integrated and separate responses in plaques and adipose tissue to weight loss and regain that have both important basic science and clinical implications.

## EXPERIMENTAL METHODS

### Animals

All procedures were done in accordance with the US Department of Agriculture Animal Welfare Act and the US Public Health Service Policy on Humane Care and Use of Laboratory Animals, and were approved by the Institutional Animal Care and Use Committee of the New York University School of Medicine (protocol number 160725-01). Eight-to-twelve weeks-old male *Ldlr^-/-^*, WT C57BL/6J (CD45.2+) and CD45.1+ mice were housed in a temperature-controlled (22^0^C) room on a 12 h light/dark cycle. Original breeding pairs were purchased from Jackson Laboratory (Bar Harbor, ME).

### Weight cycling model

Obesity was induced by feeding *Ldlr^-/-^* or WT mice a high fat-high cholesterol (HFHC) diet (Research Diets, 17052507i; New Brunswick, NJ) *ad libitum* for 24 weeks (baseline; BL). To induce LDLr deficiency and hypercholesterolemia in WT mice, they received weekly injections of 5mg/kg antisense oligonucleotide (ASO) targeting the LDL receptor (generously provided by Ionis Pharmaceuticals)(27)for the duration of HFHC diet feeding. Weight loss was induced by restricting daily caloric intake of obese mice by 30% for 2 weeks (short-term caloric restriction, stCR), as described by Ferrante and colleagues(28). After weight loss, a group of mice was again provided with HFHC diet *ad libitum* for 6 weeks (weight regain, WR).

### Glucose tolerance test

Glucose tolerance test (GTT) was performed after fasting for 6h and acclimation to the testing room with access to water. Mice were injected intraperitoneally (i.p.) with either D-glucose (Crystalgen 300-341-1000) at 2g per kg of body weight. Blood glucose levels were measured via tail sampling at baseline (t = 0) and again at t = 15, 30, 60 and 90 minutes post glucose injections, using a glucometer (Contour Next EZ, Bayer).

### Lipid and lipoprotein analyses

Total cholesterol (TC) was measured using the enzymatic assay Total Cholesterol E Kit (Wako Life Science, NC9138103; Richmond, VA). Plasma lipoproteins were separated using fast-performance liquid chromatography with two Superose 6 10/300 GL columns (GE Healthcare; Boston, MA) on a Shimadzu HPLC system (Columbia, MD).

### Plaque morphometrics and immunohistochemistry

Mice were euthanized at the end of each dietary regimen and blood was collected via cardiac puncture, followed by perfusion with saline at physiologic pressure. Hearts were then removed and embedded in OCT (Sakura, 4583; Torrance, CA) and immediately frozen at -80^0^C, while arches were removed and kept in PBS for same day flow cytometry. Aortic root sections (6 µm) were stained for CD68 (Bio-Rad MCA1957; Hercules, CA) to detect macrophages. Picrosirius red (PolySciences 24901-500; Niles, IL) was used for collagen staining, and imaging was done using polarizing light microscopy. Necrotic core was quantified by measuring hematoxylin and eosin-negative acellular areas in the intima, as described previously (29).

TUNEL staining was performed according to the manufacturer’s instructions (Invitrogen C10619; Waltham, MA). In brief, thawed aortic root sections were fixed with 4% paraformaldehyde for 15 minutes at 37^0^C and then proteinase K solution was added for 5 minutes. Slides were then washed and fixed once more with 4% paraformaldehyde for 5 minutes at 37^0^C. TdT buffer was then added for 10 minutes, removed and TdT reaction solution added for 60 minutes, after which slides were washed and incubated with 0.1% TritonX-100 for 5 minutes. Slides were then added the TUNEL reaction cocktail for 30 minutes at 37^0^C, washed and then proceeded to Fcgr4 and CD68 staining. Antibodies for both Fcgr4 (SinoBio 50036-T24; Wayne, PA) and CD68 (as above) were added at a 1:250 dilution and incubated for 1 hour at room temperature. Secondary antibodies were used at a 1:400 dilution, and added after washing with PBS. After 1 hour incubation, slides were washed again and mounted using Prolong Gold (Invitrogen P36934; Waltham, MA).

### Isolation of immune cells from aortic arch and adipose for flow cytometry

Cell suspensions from plaques were generated after aortic arch (see below) and adipose digestion(13), and sorted using BD FACS Aria II SORP. To single-cell suspensions was added a live/dead cell staining (Invitrogen, L10119; Waltham, MA) and the following antibodies:BV510 anti-CD45 (103137), PE anti-CCR2 (150609), PE-Cy7 anti-Ly6G (127618), BV711 anti-CD11b (101241), BV570 anti-Ly6C (128030), BV650 anti-CD11c (117339), BV421 anti-B220 (117339), BV421 anti-CD3 (100227), AF647 anti-CD206 (141712), PE-Cy7 anti-CD14 (123316), PerCP-Cy5.5 anti-FCGR4 (149518). All antibodies were purchased from Biolegend (San Diego, CA), as well as BV421 anti-SiglecF (BD Bioscience, 562681; Franklin Lakes, NJ) and PE anti-CD163 (eBioscience, 12-1631-80; San Diego, CA).

Aortic digestion was performed by mincing the aortic arch and branches with a blade. Minced tissues were suspended in 3mL digestion buffer: Hank’s Balanced Salt Solution, 1mM EDTA, 1% BSA, 0.77mg/mL Liberase (Roche, 273582; Basel, Switzerland), 0.1mg/mL Hyaluronidase (Sigma Aldrich, 3506; St. Louis, MO) and 0.06mg/mL DNaseI (Sigma Aldrich, DN25; St. Louis, MO). Tissue was then transferred to C-tubes, placed in a gentleMACS dissociator (both from Miltenyi Biotech; Bergisch Gladbach, Germany) and digested for 15 minutes in 37^0^C. Suspensions were filtered using a 100μm nylon mesh and centrifuged for 5 minutes. Cell pellets contained single cells that were next stained for flow-cytometry analysis.

### RNA extraction and quantitative real-time PCR analysis

Total RNA was extracted from cells using TRIzol (Invitrogen) and Direct-zol Miniprep kit (Zymo Research) followed by cDNA synthesis using the Verso cDNA kit (Thermo Scientific). SYBR Green-based real-time qPCR was performed using the ABI PRISM 7300 sequence detection system (Applied Biosystems). Gene expression was normalized to *Hprt* expression and assessed using Δ(ΔCt). Primer sequences were as follows: *Tnfa*, (Fw *5’-GATCTCAAAGACAACCAACATGTG-3’, Rv 5’- CTCCAGCTGGAAGACTCCTCC CAG-3’), Nos2, (Fw 5’- CTGATGGCAGACTACAAAGACG-3’, Rv TGGCGGAGAGCATTTTTGAC-3’)*

*IL12 (Fw 5’ TGGTTTGCCATCGTTTTGCTG -3’, Rv 5’- ACAGGTGAGGTTCACTGTTTCT-3’) Mrc1 (Fw 5’- CTCTGTTCAGCTATTGGAGCG-3’, Rv 5’-CGGAATTTCTGGGATTCAGCTTC-3’)*

*Arg1 (Fw 5’-CTCCAAGCCAAAGTCCTTAGAG-3’, Rv 5’- AGGAGCTGTCATTAGGGACATC-3’)*

### Efferocytosis

Jurkat cells were irradiated with ultraviolet light for 15 minutes to induce apoptosis, followed by labeling with CellTracker™ Green CMFDA dye (Invitrogen, C7025), for 30 min. Cells were washed once with 10 mL of serum free media and suspended in RPMI+ 10% FBS and 1% penicillin/streptomycin and kept in the incubator for 3 hours. After that, apoptotic Jurkat cells were applied to macrophages (2.5:1 Jurkat:macrophage) that had been seeded in chamber slides the previous day. After 40 min incubation, media samples were aspirated and macrophages were washed three times with PBS, followed by fixation with formalin for 15 min. Macrophages were then stained with DAPI and APC anti-CD11b. Efferocytotic events were calculated as CD11b+ cells (i.e., macrophages) with green label inside. Results were expressed as % of total macrophages.

### Isolation and treatment of primary macrophages

Macrophages were obtained from adult WT and *Fcgr4* conditional knockout (*Fcgr4^fl/f^*^l^-LysM^Cre^) mice from bone marrow and peritoneal exudate cells (PECs). Peritoneal cells were obtained by washing the peritoneal cavity with 5mL of sterile PBS. Erythrocytes were removed by incubating the cell pellet with red blood cell lysis buffer (Sigma-Aldrich, R7757; St. Louis, MO) for 5 minutes at room temperature. After washing with PBS, cells were cultured in DMEM (1gL/glucose, Corning, MT10014CM; Corning, NY) containing 10% FBS (Gemini bio-products; Sacramento, CA), 1% penicillin/streptomycin and 10ng/mL M-CSF (Peprotech, 315-02; Cranbury, NJ). Cells were seeded in chamber slides (Thermo Fisher, 154534; Waltham, MA) at 500,000 cells per well and kept in the incubator overnight.

Bone marrow derived macrophages (BMDMs) were obtained by harvesting femur and tibia, cutting the edges of each bone and spinning them for 20 seconds at 8000 rpm, in an Eppendorf tube containing 100 uL of PBS. After red blood cell lysis (as above), cells were plated in 48 well plates (40 wells per mice) in DMEM (1gL/glucose) containing 10% FBS, 1% penicillin/streptomycin and 10ng/mL M-CSF (as above). Fresh medium was added on day 3. Cells were treated with either 10ng/mL LPS (; Sigma Aldrich, L4391; St. Louis, MO) or IL-4 (Peprotech, 214-14; Cranbury, NJ) for 6 hours, or re-seeded in chamber slides (500,000 cells per well, as above) and incubated overnight for the efferocytosis assay (see above) on the following day.

Total bone marrow cells were obtained by flushing bone marrow from femurs and tibias and plating on 96-well flat-bottom well culture plates (270,000 cells/well) in complete DMEM, as above. After a 2-hour resting period in the incubator (37^0^C, 5% CO_2_), cells were stimulated for 16 hours with 100ng/mL LPS, or media only (as control). Subsequently, supernatants were collected for cytokines measurement through ELISA.

### Bone marrow progenitor isolation and quantification

Bone marrow cells were obtained from femur and tibia (same as for obtaining BMDMs, as described above). After the red blood cell lysis, cells were counted and added a green live/dead stain (Invitrogen, L34969; Waltham, MA) in PBS and kept at 4°C, in the dark, for 30 minutes. Cells were then spun, supernatants discarded, and to the cells was added a cocktail of antibodies: from eBioscience (San Diego, CA)-FITC anti-GR1 (11-5931-82), FITC anti-CD3 (11-0031-82), FITC anti-CD4 (11-0041-82), FITC anti-CD8 (11-0081-82), FITC anti-Ter119 (11-5921-82), FITC anti-CD19 (11-0193-82), FITC anti-NK1.1 (11-5941-82), FITC anf-CD2 (11-0021-82), APC anti-CD34 (50-0341-82), and from Biolegend (San Diego, CA)-FITC anti-CD11b (101206), PE-Cy7 anti-Sca1 (108114), APC-Cy7 anti-cKit (105826), PE anti-CD135 (135306), PerCP-Cy5.5 anti-CD150 (115922), BV605 anti-CD48 (103441) and BV711 anti-CD16/32 (101337). Cells were kept at 4°C, in the dark, overnight, after which they were washed and analyzed using an LSRII (BD Biosciences; Franklin Lakes, NJ).

### Adipose transplantation

CD45.1 adipose tissue donors were fed a HFHC diet for 24 weeks, after which some mice were calorically restricted for 2 weeks (given 70% of their daily food consumption, as above). Recipient mice were lean male *Ldlr^-/-^* with established atherosclerosis, achieved by low-fat high-cholesterol diet feeding (Research Diets)(30) for 20 weeks. The use of a low-fat diet prevented the potentially confounding development of obesity. The transplantation studies were conducted following standard antiseptic surgical techniques. eWAT from either obese or stCR CD45.1 mice was immediately excised post-euthanasia, washed in sterile PBS, and 400mg were transplanted into the subcutaneous anterior dorsal region of recipient mice, as previously described(31). Two days pre-transplant, recipients were switched to a low-cholesterol diet (chow), to gradually lower plasma cholesterol and allow reparative signals (if any) from the transplanted adipose tissue influence plaques or bone marrow. Tissues of eWAT recipients were harvested 2 weeks post-transplant.

### Murine white blood cell (WBC) counts

Total WBC counts in freshly isolated blood were performed by collecting tail blood in EDTA-contained tubes and analyzed using a hematology cell counter (Heska Element HT-5).

### *Fcgr4* siRNA particles and treatments

Particles containing scrambled (Dharmacon D-001810-04; Lafayette, CO) or *Fcgr4* (Dharmacon J-053664-17) siRNA were generated as previously described(32–34). WT males were injected with *Pcsk9* AAV.8TBGmPCSK9D377Y (2 × 10^12^ viral particles/mouse, Penn Vector Core; Philadelphia, PA), and placed on a HFHC diet to induce LDLr-deficiency and atherosclerosis. After 20 weeks the obese mice were randomized to groups with similar body weight and stCR was started. At the same time mice were injected with 200uL particles i.p. once a day for 14 days.

### Bone marrow (BM) transplantation

BM cells were harvested from femora and tibias of *Ldlr^-/-^* donor mice (BL, CR, WR). After red blood cell lysis (as above), 1x10^6^ cells were suspended in 0.2 mL PBS and injected retro-orbitally into male CD45.1 recipients that were lethally x-irradiated with a total dose of 10Gy. After 4 weeks of recovery, recipient mice were injected i.p. with the *Pcsk9* AAV8 (as above). Animals were provided with HFHC diet for 14 weeks and harvested thereafter.

### Cytokine ELISA

Purified and biotinylated IL-6 (504502 and 504601, respectively) and IL-10 (505002 and 504906, respectively) antibodies were purchased from Biolegend (San Diego, CA). Purified antibodies were used to coat an ELISA plate, according to the manufacturer’s instructions, and incubated overnight in 4^0^C. The plate was then washed 4 times in PBS containing 0.05% tween (PBST) and blocked with 1% BSA in PBS. After 1 hour incubation at room temperature, medium conditioned by bone marrow cells was added and incubated in 4^0^C overnight. The plate was then washed 4 times in PBST and added was one of the biotinylated antibodies, according to the manufacturer’s instructions, and incubated for 1 hour at room temperature. The plate was then washed 4 times in PBST and next added was alkaline phosphatase conjugated to streptavidin, from Jackson ImmunoResearch (016050084; West Grove, PA) according to the manufacturer’s instructions. After 30 minutes incubation at room temperature the plate was washed 4 times with PBST and alkaline phosphatase substrate was added (Sigma Aldrich, S0942; St. Louis, MO). The plate was analyzed using a plate reader, for emission in 405nm. Standard curves were established using mouse recombinant IL-6 (Peprotech, 216-16; Cranbury, NJ) and IL-10 (Biolegend, 575804; San Diego, CA).

### Methods for scRNAseq analysis

#### Read alignment, barcode de-convolution, and UMI (unique molecular identifier) counting

Following sequencing, we used the CellRanger Single Cell Software Suite v 3.1.0 to de-multiplex individual cells, process UMIs, and count UMIs per gene, following the standard pipeline and default parameters. Briefly, by using *cellranger count*, FASTQ files were generated and aligned to the mouse mm10 genome, sequencing reads were filtered by quality score, and cell barcodes and UMIs were assigned to each read. The filtered gene expression matrices from the plaque BL and stCR samples were then used for downstream analyses.

#### Filtering cells

To identify low-quality cells and doublets, we visualized the distribution of the number of UMI detected for each cell, the number of genes expressed, and the percentage of reads that were coming from mitochondrial genes using Seurat v 4.1.1(35). We removed cells that were high outliers for the number of UMI or the number of genes, as these may reflect doublets, i.e. multiple cells captured in a single gel bead in emulsion (GEM). We also removed cells that were high outliers for mitochondrial gene expression, indicating cells undergoing apoptosis.

#### Normalization and alignment of datasets

To merge our plaque datasets with those from Weinstock et al.(13), we first normalized each of the 4 samples (Plaque BL and CR, and Fat BL and CR) using the *SCTransform* within Seurat. We then integrated the datasets using *SelectIntegrationFeatures* with default parameters and *nfeatures =3000*. We then ran *PrepSCTIntegration*, *FindIntegrationAnchors* and *IntegrateData* using default parameters, but integrating all genes in the dataset, not just the integration features.

#### Dimensionality reduction, cluster finding, and cell type annotation

After integrating the 4 datasets, we ran the following functions on the “integrated” assay using Seurat: *RunPCA*, *RunUMAP*, *FindNeighbors*, and *FindClusters*, using 30 PCAs for *RunUMAP* and *FindNeighbors*. We then annotated the clusters with SingleR 1.4.1, using the dataset from the Zernecke et al. (2020) meta-analysis as a reference dataset.

#### Downsampling and differential expression

The plaque BL sample had significantly deeper sequencing coverage than the other 3 samples, as shown in Supp. Fig. 2A. This resulted in a skew towards genes that had higher expression in plaque BL when we ran differential expression analyses. To correct this likely source of bias, we downsampled all 4 samples to include a maximum of 10,000 UMI per cell using the *SampleUMI* function within Seurat. We retained the same cluster identities as in the full dataset, and ran differential expression using the downsampled data to get more symmetrical numbers of differentially expressed genes between Plaque BL and the other cell types.

#### Cell-cell communication

Metadata information on cluster membership and raw counts were extracted for each sample and normalized as counts per 10,000; these were then used to run the *cellphonedb method statistical_analysis*(36) for each individual sample for the *within-tissue* analysis. The *significant_means* output was then parsed using custom scripts to plot the total number of significant interactions per cluster pair. For the *cross-tissue* analysis, data were processed in the same way except that cells from both tissues of the same condition were processed simultaneously. Again, the *significant_means* output was queried, with ligand-receptor identified between cluster pairs within the same tissue removed.

#### Bulk RNA sequencing analysis

Samples from the bulk sequencing analysis were first mapped to mm10 using STAR v 2.6.0, and gene counts were generated using the FeatureCounts function within subread v 1.6.3. We then used DESeq2 v 1.30.1 to detect differential expression between the Fcgr4 positive and Fcgr4 negative macrophages, using a paired sample approach with the model *design = ∼mouse + treatment* where mouse was the individual mouse and treatment was either Fcgr4 positive or negative. We used the Bioconductor package clusterProfiler v 3.18.1 to perform GO and KEGG enrichment on genes significantly differentially expressed between Fcgr4 positive and negative macrophages.

### Data availability

Data are publicly available in GSE225077 (bulk RNA-seq of Fcgr4+ macrophages) and GSE141036 (scRNA-seq of eWAT and plaque leukocytes).

## RESULTS

### Short-term caloric restriction in obese mice promotes atherosclerosis resolution

We previously showed in multiple mouse models with hypercholesterolemia that substantial lipid lowering results in the resolution of atherosclerotic plaques as judged by decreased content and inflammatory properties of macrophages(24–26). In order to isolate effects of weight loss in obesity on established atherosclerosis, a model in which cholesterol levels are not dramatically affected was required. Another consideration in study design was that switching the feeding of a high-fat diet to normal chow results in a severe reduction in food intake(37) (mimicking long-term fasting), as well as in epigenetic changes in macrophages and their precursors related to differences in diet composition(15, 16). Thus, we adapted a protocol of mild(28) short term caloric restriction (stCR), keeping the diet composition the same, in order to investigate the role of clinically relevant reduced caloric intake and weight loss on inflammation resolution in atherosclerosis independent of cholesterol lowering.

Thus, LDLr-deficient mice(27) were fed a high-fat high-cholesterol (HFHC) diet *ad libitum* for 24 weeks to develop obesity and advanced atherosclerotic plaques. At this time, tissues from the baseline (BL) group were harvested. To examine early changes induced by weight loss, the stCR group was switched to daily feeding of 70% of their *ad libitum* consumption of the same HFHC diet for an additional 2 weeks (Fig. 1A). The data show that after 24 weeks of treatment, mice were obese, and after 2 weeks of stCR, they lost 14.3% of their weight (respective mean weights of 44.9±1.37g and 38.5±1.6g; Supp. Fig. 1A). Upon harvest, several tissues were weighed and a reduction in eWAT mass was observed (Supp. Fig. 1B) with no significant changes to the masses of inguinal white AT (iWAT), brown AT (BAT), liver or kidney (Supp. Fig. 1C-1F).

**Fig. 1:**
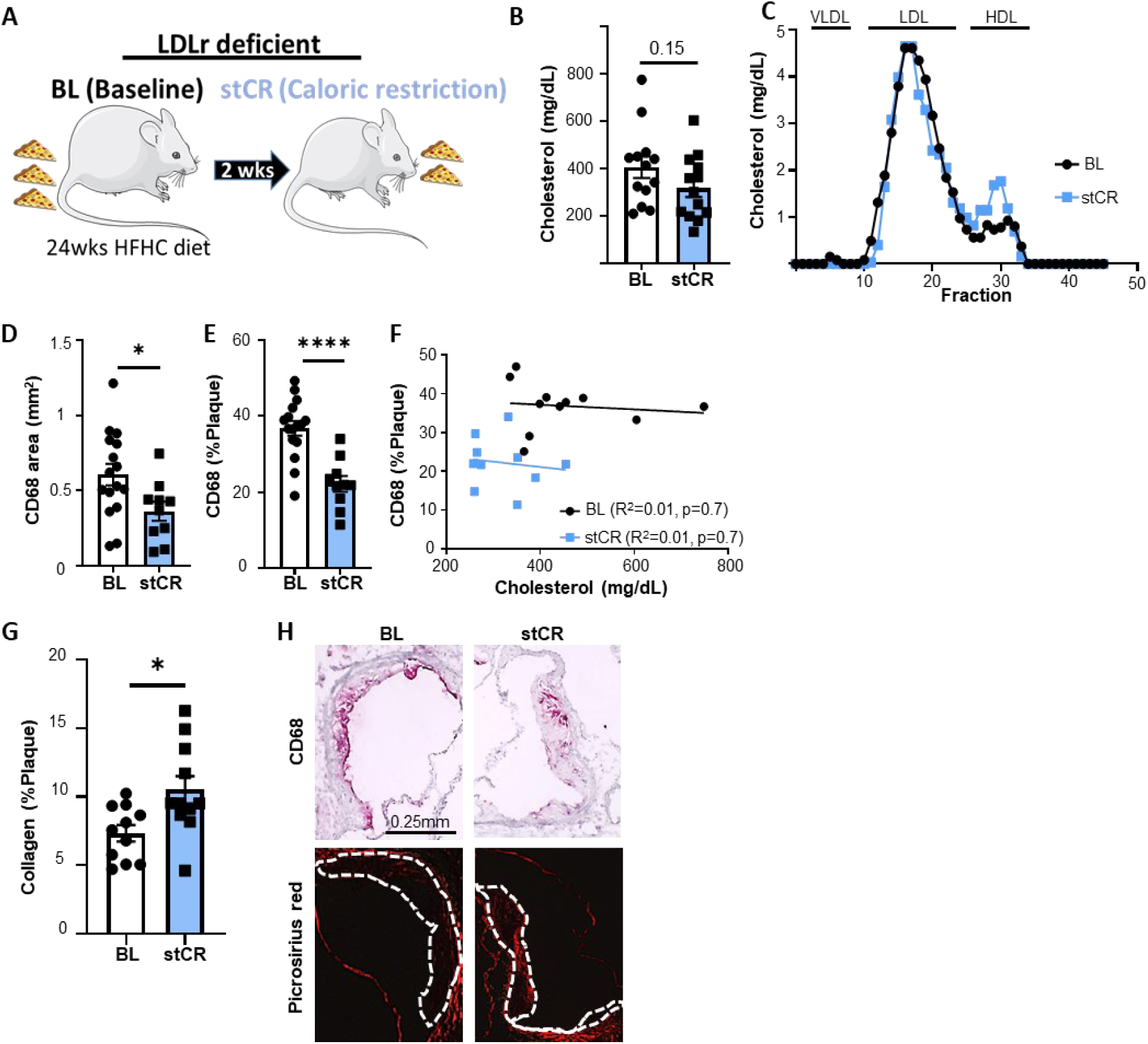
stCR induces atherosclerosis resolution. (A) Experimental design. Mice were fed a high-fat, high cholesterol (HFHC) diet for 24 weeks to induce obesity. Mice were then calorically restricted for 2 weeks (stCR), through reducing daily food intake by 30%. (B) Plasma cholesterol levels and (C) lipoprotein profile at the end of the experiment. (D, E) Plaque macrophage content quantified through CD68 staining. (F) Simple linear regression showing lack of correlation between cholesterol levels and CD68 content in plaques. (G) Collagen quantification in plaques assessed through Picrosirius red staining. (H) Representative aortic root images. **\***p<0.05, ****p<0.0001 determined via two-tailed Student’s t-test.

Examination of metabolic parameters showed marked improvements with stCR, including reduced fasting glucose (Supp. Fig. 1G), lower HOMA-IR (a measure of insulin resistance; Supp. Fig. 1H) and improved glucose tolerance (Supp. Fig. 1I-J). Moreover, plasma cholesterol levels remained elevated after stCR, with non-significant changes between the two groups (Fig. 1B). To investigate the lipoprotein distribution of plasma cholesterol, plasma samples were fractioned by Fast Protein Liquid Chromatography (FPLC) and cholesterol was measured in each fraction. The results showed no significant differences in the distributions between the groups (Fig. 1C). For the evaluation of atherosclerosis, aortic roots were sectioned and investigated for plaque size and composition. While plaque area was comparable between the groups (164,424±18,802µm^2^ in BL and 149,934±26,641µm^2^ in stCR; Supp. Fig. 1K), the stCR group had fewer macrophages, observed both as a decrease in the area of CD68+ cells (Fig. 1D) and their proportion of the total plaque area (Fig. 1E). In an independent analysis, consistent findings were found from digested aortic arches that were analyzed using flow-cytometry. These results showed fewer macrophages in aortic arches of stCR mice, compared to the BL group (Supp. Fig. 1L). To further establish that the changes in the macrophage contents of atherosclerotic plaques were independent of plasma cholesterol levels, we investigated whether these parameters were correlated. Statistical analysis shown in Fig. 1F demonstrated no correlation between the two parameters. The change in plaque cellular composition without a decrease in area is reminiscent of some of our previous studies (e.g.,(26)), in which inflammation-resolving plaques also did not reduce in area. Rather, they become enriched in collagen, presumably because the content of matrix metalloprotease-producing (inflammatory) macrophages decline. In human plaques, such enrichment is taken as a sign of increased stability(38). To quantify changes in the collagen content of plaques, aortic root sections were stained with picrosirius red and the positive areas were quantified from polarized light images. Indeed, consistent with our previous data, concurrent with decreased plaque macrophages, there was increased collagen content following stCR (Fig. 1G). Representative images of aortic roots stained for CD68 and picrosirius red are presented in Fig. 1H.

### Macrophages play a major role in the inter and intra-organ communication in obesity and stCR

To investigate at the molecular level how stCR influences the immune compartment in atherosclerotic plaques, first, single-cell suspensions were obtained from aortic arches harvested from mice in both experimental groups. Viable CD45+ cells (i.e., all leukocytes) were sorted and transcripts of individual cells were sequenced, using the 10× Genomics platform (following the method described in(39)). Because we have also obtained adipose tissue CD45^+^ single-cell transcriptomic data from the same mice(13), the gene expression profiles from both tissues were merged. Quality control and data filtering are displayed in Supp. Fig. 2A.

Unbiased clustering found 23 distinct clusters (Fig. 2A, Supp. Fig. 2B). To annotate the different clusters, we used a recently published meta-analysis of plaque single-cell transcriptomes as a reference dataset(40) (Supp. Fig. 2C-D). The top 5 differentially expressed genes (DEG) in each cluster are presented in Supp. Fig. 2E. Many of the clusters in our dataset corresponded with previously published work(40); however, some unique clusters were found as well. For these, we used our previously published dataset from the eWAT CD45^+^ cells (Fig. 2A) as the reference dataset(13). Cell proportions were plotted for each tissue in BL and stCR conditions (Fig. 2B). Note that while all clusters are shared across eWAT and plaques, their distributions considerably differ in both the obese and stCR conditions. Similar to previous reports(40, 41), we also found that macrophages had the greatest heterogeneity in both eWAT and plaques, constituting 6 of the clusters.

**Fig. 2:**
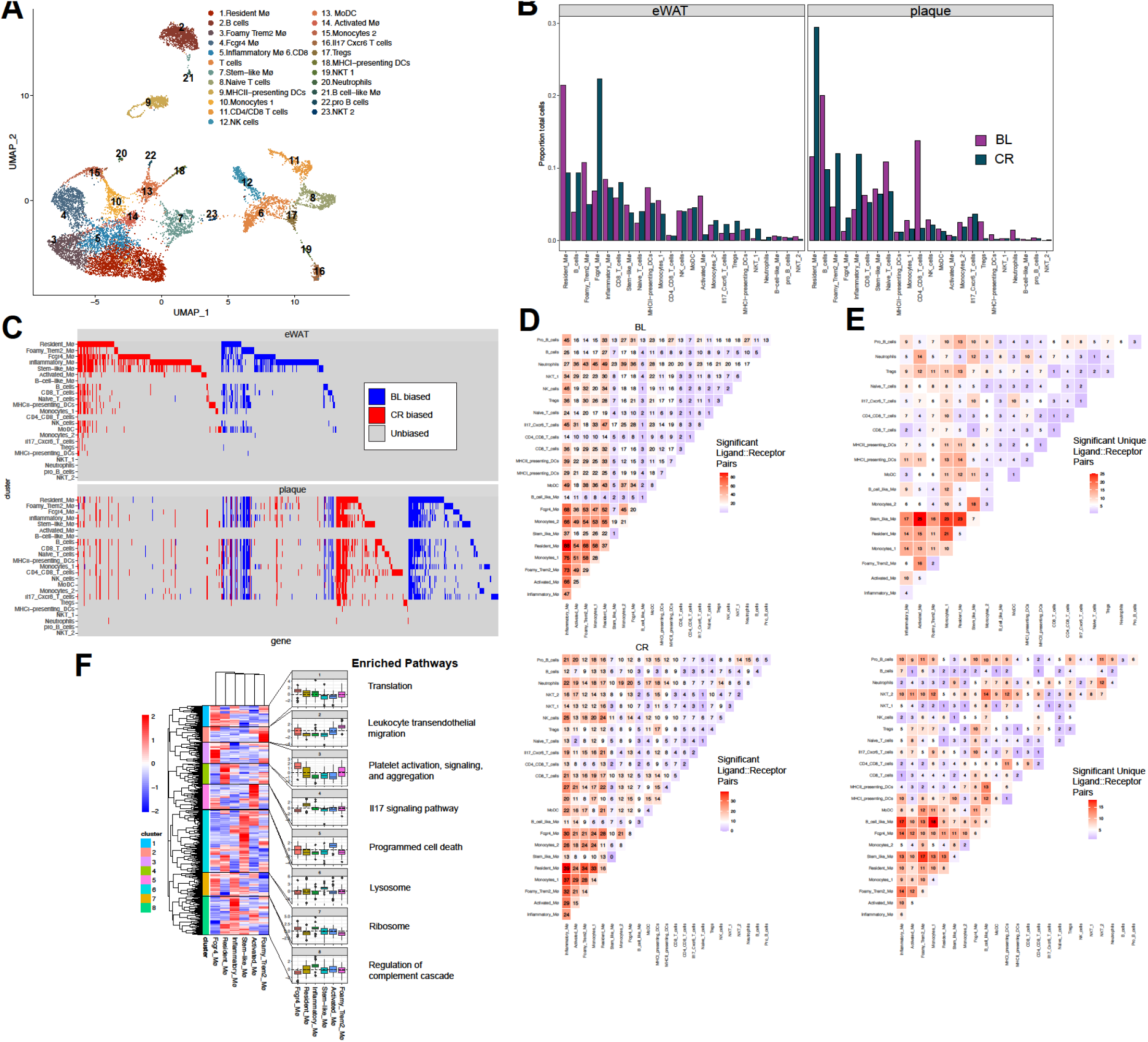
Inter and intra organ communication of immune cells in obesity and weight loss. (A) Unbiased clustering and cell type assignment of scRNAseq of CD45+ cells from eWAT and aortic arch represented in an UMAP. (B) Proportion of each cell cluster identified in the scRNAseq analysis by tissue of origin. (C) Genes (columns) differentially expressed in stCR compared to BL are plotted per cluster (rows) in eWAT (top) and plaque (bottom). In blue and red are genes that are significantly upregulated in BL and stCR, respectively, or unchanged in grey. (D) Cell-cell interaction analysis, representing the number of receptor-ligand pairs across all clusters in the plaque of BL (top) and stCR (bottom) conditions. (E) Receptor-ligand interaction analysis across tissues in BL (top) and stCR (bottom), after subtracting pairs observed in within-tissue interactions. (F) Hierarchical clustering of DEGs between stCR and BL in all macrophage clusters of the plaque scRNAseq dataset, and their associated enriched pathways.

We also investigated whether obesity and stCR drove similar gene expression in *both* eWAT and plaques, as well as in distinct leukocyte clusters within each tissue. The expression of each DEG in eWAT was plotted per leukocyte cluster, and its corresponding expression in plaques is shown in Fig. 2C. We classified all DEGs (columns) as either BL-biased, with statistically significant higher expression in BL (blue), or stCR-biased, with higher expression in stCR (red) across all clusters (rows). Many columns (i.e., DEGs) show a signal in multiple rows (cell clusters), indicating that several clusters differentially express the same genes within each tissue, and often in the same direction (either BL or stCR-biased). To look further into this, we plotted the number of clusters that shared DEGs in each tissue (agnostic of whether they are BL or CR-biased). Supp. Fig. 2F shows that although most changes in gene expression are not coordinated across clusters (indicated by the highest bars being the DEGs that are shared across 0-1 clusters), numerous genes were differentially expressed in multiple clusters, with some changing coordinately in >15 (Supp. Fig. 2F), further reinforcing that some genes change in a coordinated manner across clusters within each tissue.

We next investigated whether genes change concordantly *across* tissues (Fig. 2C). When looking at individual genes (columns) across tissues (top and bottom panels), a number appear to be coordinately regulated in eWAT and plaque (as reflected by showing signals in the same columns in the top and bottom plots). This suggests that a core set of genes (Supp. Table 1) is regulated similarly not only between clusters, but also across tissues (e.g., same column in Fig. 2C top and bottom). Most genes, however, appear to be uniquely differentially expressed in one tissue or the other (i.e., not showing concordant signal in Fig. 2C top and bottom). For instance, *Fabp4* is a gene that has increased expression in stCR compared to BL (stCR-biased) in 14 clusters from the eWAT, but none in the plaque (Supp. Table 1). *Fabp4* is an adipokine that was previously shown to stimulate insulin secretion from pancreatic β-cells (reviewed in (42)). *Fabp4* deficiency protects against diet-induced obesity(42). It is possible that *Fabp4* coordinated upregulation in the eWAT with stCR is aimed at limiting weight loss with undernutrition. Supp. Table 2 summarizes all DEGs shared by 5 or more clusters, further indicating if the expression is BL or stCR biased, and in which tissue.

Next, we aimed to infer cellular communications between the 23 cell clusters. Ligand-receptor interaction analysis was performed (see Methods), and the number of interacting pairs in plaques plotted in Fig. 2D. The corresponding interactome analysis for the eWAT is shown in Supp. Fig. 2D. The color represents the number of receptor-ligand pairs found between clusters in the X and Y axes, with the exact number of interactions indicated within each box. Most notably, in both plaque and eWAT, the cells with the highest number of significant ligand-receptor interactions among the leukocytes are the monocytes/macrophages, which mostly communicate with other macrophages (Fig. 2D, Supp. Fig. 2G and Supp. Table 3).

Using similar computational methods, we next investigated potential cross-tissue interactions of leukocytes. Ligand-receptor pairs were again examined; however, all pairs that were found in one tissue or the other were excluded, to ensure these are not artifacts of within-tissue interactions. In this way we limited ourselves to looking only at ligand-receptor pairs that are novel in the cross-tissue analysis. As a result, we found fewer interactions inter-organ compared to intra-organ. We did find, however, some potential unique means of communication, especially related to macrophages, between eWAT and plaque leukocytes (Fig 2E, Supp. Table 4). For instance, many clusters in the plaque show expression of chemokines (e.g., *Ccl2, Ccl24*), while several immune clusters in the eWAT express corresponding chemokine receptors (e.g. *Ccr2*, Supp. Table 4). Plausibly, leukocytes from adipose tissue can be recruited to plaques via such ligand-receptor interaction, which is relevant to our fat transplant experiments described below. Notably, stCR did not substantially alter the amount of intra and inter-organ interactions of leukocytes, implying that the magnitude of intra-organ communication as judged bioinformatically is not regulated by the mouse’s metabolic state (Fig. 2E, Supp. Table 4 and 5). Taken together, the foregoing bioinformatic results demonstrate that stCR drastically alters the immune landscape in eWAT and atherosclerotic plaques, and changes gene expression concordantly in most clusters.

To further explore the responses of each macrophage cluster to stCR, the log fold change (LFC) values compared to BL were hierarchically clustered and examined for pathway enrichment (Fig. 2F). First, we performed differential expression analysis (see Methods) between stCR and BL within each macrophage cluster, and then calculated LFC values for those genes in all other macrophage clusters. We then performed hierarchical clustering on the LFC values across genes (rows) and macrophage clusters (columns).

There were 8 well-defined clusters capturing distinct patterns of differential gene expression across the macrophage clusters. The differential expression patterns of genes in each hierarchical cluster were summarized using boxplots to surmise the LFC patterns across clusters. GO and KEGG analyses to assess pathway enrichment in each hierarchical cluster were then performed, and the resulting terms are shown in Fig. 2F and Supp. Table 6.

Notably, the hierarchical clustering showed that the Fcgr4+ macrophage cluster has the most distinct response to stCR, as it is the basal group in the hierarchical tree (Fig 2F). Interestingly, we have previously observed these cells to accumulate in the eWAT in obese mice following stCR, and they express many genes associated with phagocytosis(13), a process that would be expected to have important functional effects in both tissues. Therefore, we chose to investigate these cells further, and examine whether they play a role in stCR-induced atherosclerotic plaque resolution.

### Fcgr4+ macrophages accumulate with weight loss and promote beneficial changes in atherosclerotic plaques

As just noted, in both plaques and eWAT, stCR strikingly enriched macrophages distinguished by high expression of *Fcgr4* (Fig. 2A and 3A). These transcriptomic results were verified using immuno-fluorescent staining of aortic roots and eWAT sections for macrophages (CD68 and F4/80, respectively) and FCGR4 (Fig. 3B).

**Fig. 3:**
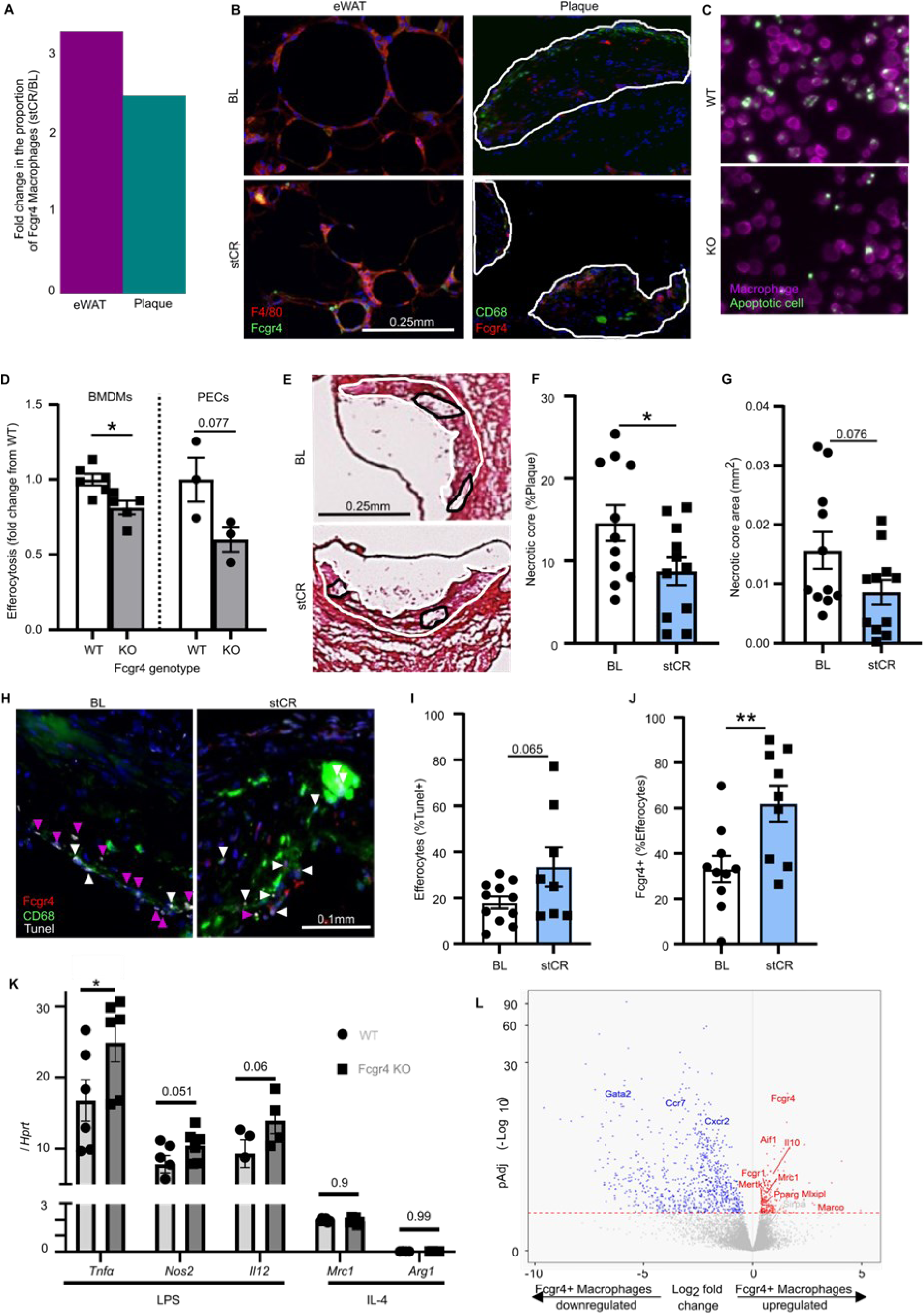
Fcgr4+ macrophages accumulate with stCR and promote a pro-reparative phenotype through increased efferocytosis. (A) Fold change in the proportion of Fcgr4+ macrophages in stCR mice, compared to baseline, in plaque and adipose tissue, quantified from the scRNA-seq data. (B) Representative images of FCGR4 and macrophage staining in eWAT and aortic roots. (C, D) BMDMs and PECs from WT and *Fcgr4* KO mice were exposed to fluorescent labeled (green) apoptotic Jurkat cells. Efferocytotic events were determined as macrophages having an attached or engulfed green label. (E-G) Plaque necrotic core quantification in root sections of baseline and stCR mice. (H) In situ efferocytosis assay of aortic root sections in which apoptotic cells were labeled by TUNEL (white), macrophages by anti-CD68 (green), FCGR4 (red) and nuclei by DAPI (blue). White arrows indicate macrophage-associated TUNEL and purple arrows mark free TUNEL. Efferocytosis was calculated as (I) total efferocytes (TUNEL+ macrophages) and as (J) FCGR4+, TUNEL+ macrophages (J). (K) Gene expression of inflammatory (*Tnfa, Nos2, and Il12*) and anti-inflammatory (*Mrc1 and Arg1*) markers following LPS or IL-4 stimulation of WT or *Fcgr4* KO for 16 hours. (L) Volcano plot showing DE genes in Fcgr4+ macrophages from adipose tissue following stCR. **\***p<0.05, **p<0.01 determined via two-tailed Student’s t-test.

We hypothesized that the functional consequence of the enrichment would be enhanced tissue repair (specifically, inflammation resolution and favorable tissue remodeling), as suggested by Fc-receptors being potent mediators of phagocytosis (reviewed in(17)). Moreover, these macrophages were found to be enriched in other phagocytosis-related genes(13). There is strong rationale for this hypothesis from the recognition in the atherosclerosis field that efferocytosis, or the phagocytosis of dying cells, is an inflammation resolving process that limits the size of plaque necrotic cores(18). Thus, we investigated the efferocytotic function of Fcgr4+ cells in bone marrow-derived and peritoneal macrophages isolated from WT and *Fcgr4* conditional knockout mice. In a standard assay(29), they were exposed to fluorescent apoptotic cells, with efferocytotic activity quantified by how many macrophages took up these cells. As shown in Fig. 3C and D, both peritoneal and bone marrow-derived *Fcgr4*^-/-^ macrophages had reduced efferocytotic capacity compared to *Fcgr4*-sufficient cells. To examine whether stCR induces efferocytosis *in vivo*, plaque necrotic core size was assessed in aortic root images, which, as implied above, has been shown to inversely correlate with macrophage efferocytotic activity(29, 43). Indeed, the data show smaller necrotic cores in the stCR group (Fig. 3E-G).

To further examine macrophage efferocytotic capacity *in vivo*, plaques were also assessed (Fig. 3H-J), as described previously(29), for macrophage-associated apoptotic cells (observed as TUNEL+). The results show that compared to obese BL mice, plaques from the stCR group tended to have more macrophages associated with apoptotic cells (Fig. 3H-I), again, indicative of increased efferocytosis. Notably, we previously reported in eWAT that after stCR, the content of macrophages that had multiple nuclei increased, consistent with their efferocytosis of apoptotic cells(13). Here we found that in both BL and stCR mice a sizeable proportion of the efferocytes in plaques were Fcgr4+, with substantially more efferocytes expressing Fcgr4 in the stCR group (33% in BL and 62% in stCR plaques; Fig. 3J).

We also investigated whether stCR influences the expression of inflammatory and pro-resolving genes by performing flow-cytometric analysis of plaque macrophages (Supp. Fig. 3A). This revealed that stCR increased levels of proteins associated with pro-resolution (e.g., CD163 and CD206) and decreased levels of those associated with inflammation (e.g., Ly6C and CD14). Furthermore, when exposed to the inflammatory stimulus lipopolysaccharide (LPS) *in vitro*, *Fcgr4^-/-^* bone marrow-derived macrophages showed increased expression levels of pro-inflammatory genes, including *Tnfa*, *Nos2* and *Il12*, without changing the expression levels of pro-resolving genes in response to IL-4 (Fig. 3K).

We sought to further characterize Fcgr4+ macrophages by performing bulk RNA-sequencing on eWAT macrophages that were either Fcgr4+ or negative. For that, obese mice were subjected to stCR, after which eWAT was isolated, digested to single cells and Fcgr4 positive and negative macrophages were flow-sorted. DEG expression between macrophages expressing Fcgr4 and those not expressing it is presented in Supp. Table 7. As seen in Figure 3L, we found many genes that were statistically significantly upregulated in the Fcgr4+ macrophages and are known to be important in the efferocytosis process, such as *Mertk*(44)*, Il10*(45) and *Pparg*(46), further adding evidence of their phagocytic/efferocytic function. We also queried (Fig. 3L) the statistically significantly up and down-regulated genes of Fcgr4+ macrophages for KEGG pathway enrichment (Supp. Fig. 3B). Both “Response to interferon ɣ" and “Antigen processing and presentation" were enriched in the genes with higher expression in Fcgr4+ macrophages, while “Extracellular matrix organization”, “Regulation of angiogenesis”, and "Collagen fibril organization” were enriched in the genes with lower expression in Fcgr4+ macrophages. Interestingly, genes involved in “Cell killing” were enriched in genes both up- and down-regulated in Fcgr4+ macrophages, suggesting that different components of these pathways are at play in Fcgr4+ macrophages compared to Fcgr4-macrophages.

Taken together, these data suggest that stCR induces a desirable environment in both plaques and eWAT that is associated with decreased expression of inflammatory genes, increased expression of pro-resolving genes and enrichment of Fcgr4+ macrophages, which promote clearance of apoptotic cells.

*eWAT-derived Fcgr4+ macrophages contribute to the reduction in plaque necrotic core* The data above suggest that Fcgr4+ macrophages promote a reduction in plaque necrotic core upon stCR-induced weight loss. Despite being relatively enriched to similar levels in both plaques and eWAT following stCR (Fig. 3A), Fcgr4+ macrophages, in terms of cell frequency among all leukocytes, constituted a much larger population in eWAT (Fig. 2B)(13). Thus, we hypothesized that Fcgr4+ macrophages in eWAT may contribute to resolving atherosclerotic inflammation by inter-organ mechanisms.

To test this hypothesis, we performed adipose tissue transplantation studies, adapting a protocol we previously used(4). Experimentally, 400mg of eWAT from obese mice pre or post-stCR were transferred to lean *Ldlr*^-/-^ mice with established atherosclerosis (accomplished by low-fat high-cholesterol diet feeding for 20-weeks to avoid confounding effects of obesity, as in(30)). To investigate trafficking between adipose tissue and atherosclerotic plaques, we incorporated into the protocol eWAT donors with the congenic pan-leukocyte marker CD45.1, and as recipients, CD45.2+ *Ldlr*^-/-^ mice with atherosclerosis (Fig. 4A). Two weeks post-eWAT transplantation, aortae were harvested and examined by flow-cytometry and immunohistochemistry. Single-cell suspensions from aortic arches were analyzed for leukocyte populations that originated from the eWAT (i.e., CD45.1+).

**Fig. 4:**
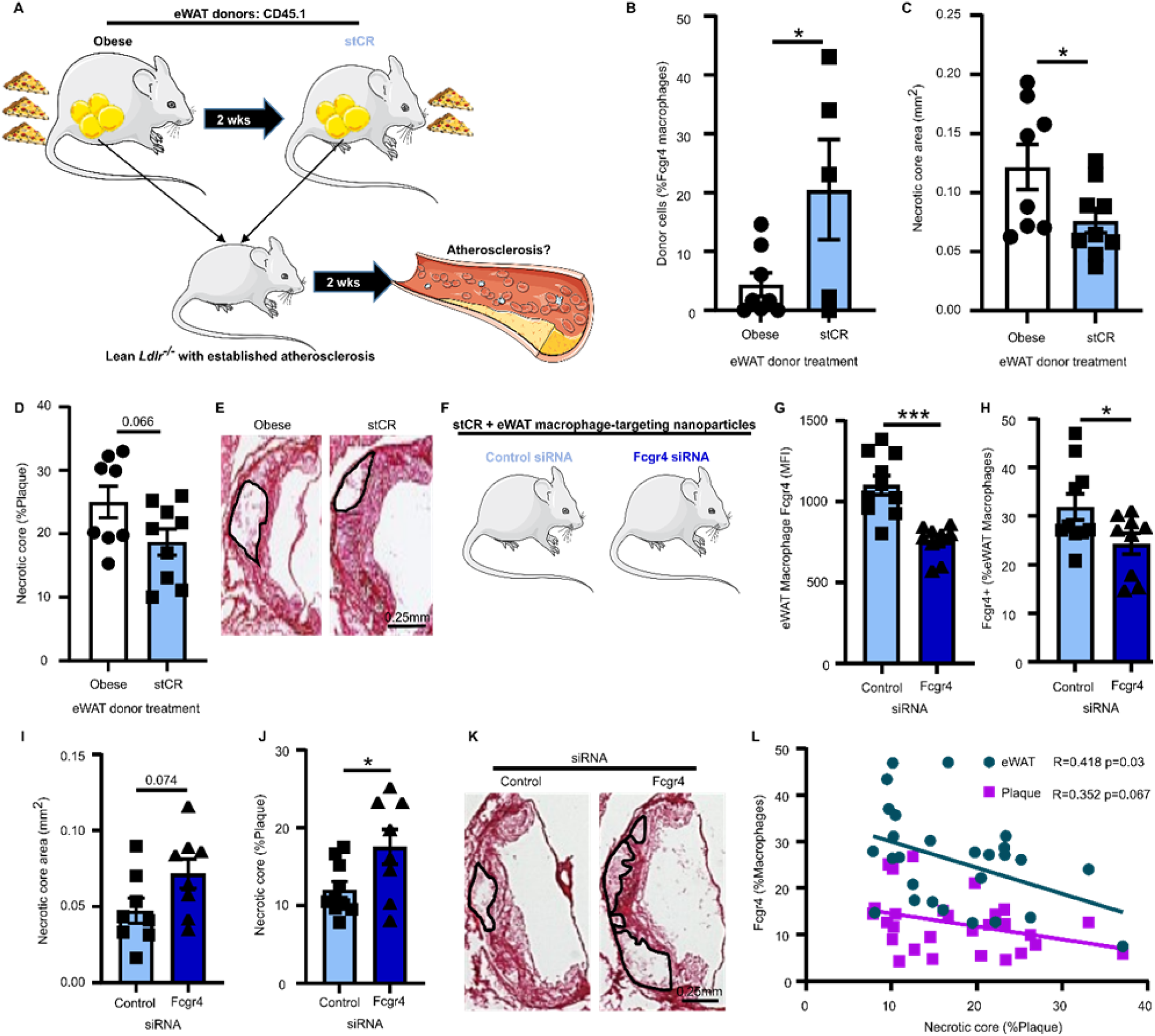
eWAT-derived Fcgr4+ macrophages reduce plaque necrotic core. (A) Schematic of adipose tissue transplantation experiment. (B) Presence of Fcgr4+ macrophages derived from donor adipose tissue in plaques of recipient mice, determined by flow cytometry. (C, D) Plaque necrotic core quantification in root sections of recipients, according to donor’s group treatment, with representative images (E). (F) Experimental design of *Fcgr4* knockdown in eWAT macrophages during CR. (G,H) Flow cytometry analysis of FCGR4 in eWAT after injection of control or *Fcgr4* siRNA particles. (I, J) Plaque necrotic core quantification in root sections, with representative images (K). (L) Simple linear regression showing correlation between Fcgr4+ macrophages and necrotic core. **\***p<0.05, ***p<0.001 determined via two-tailed Student’s t-test and (L) simple linear regression analysis.

Approximately 3% of plaque leukocytes are derived from the eWAT in recipients of either obese or stCR eWAT (Supp. Fig. 4A). Despite similar trafficking of ATMs to plaques, there was preferential enrichment of eWAT-derived Fcgr4+ macrophages in plaques of recipients of stCR eWAT (Fig. 4B). On average, 4.4% and 20.5% (p=0.04) of plaque Fcgr4+ macrophages were derived from the eWAT transplanted from obese and stCR, respectively. While there is evidence of trafficking of CD45+ cells from eWAT to plaques, it is possible that the apparent increase with stCR eWAT is driven in part by the difference in macrophage subpopulation abundances in the donor adipose tissue (Fig. 3A).

Although histological analyses of aortic root sections showed similar plaque size and macrophage content in recipients of either obese or stCR eWAT (Supp. Fig. 4B-D), plaque necrotic cores were smaller in the recipients of the stCR tissue (Fig. 4C-E). This suggested that the source of the eWAT regulates plaque necrotic core content, with a contribution related to the trafficking of Fcgr4+ macrophages. This also implies that Fcgr4 was not just a marker of cells that accumulate in eWAT with stCR, but a functional contributor to the necrotic core improvement.

To test this, we knocked down *Fcgr4* expression in eWAT macrophages during the stCR phase. For this, WT mice were injected with *Pcsk9*-AAV8 to induce LDLr deficiency and fed a HFHC diet to promote the development of both atherosclerosis and obesity. After 24 weeks, mice were randomized to have similar average weight, and stCR begun for 2 weeks. During the stCR phase, mice received daily intraperitoneal injections of particles containing siRNA to *Fcgr4*, or a scrambled sequence as control (Fig. 4F). Note that these particles are specifically taken up by macrophages in adipose tissue when obese mice are injected i.p.(32).

Pilot studies showed that the particles were taken up by 38% of eWAT and 4% of iWAT macrophages. In contrast, the particles were not detected in the spleen or bone marrow (Supp. Fig. 4E). Flow-cytometry data showed that *Fcgr4* siRNA-containing particles decreased Fcgr4 surface levels in eWAT (Fig. 4G-H), but not plaque macrophages (Supp. Fig. 4F). Of the eWAT macrophages that were positive for the particles, Fcgr4 levels were 32% higher in the control particle treated group compared to the *Fcgr4* siRNA particle-treated mice (Fig. 4G). This corresponded to a 25% decrease in the abundance of eWAT Fcgr4+ macrophages (Fig. 4H) in the *Fcgr4* siRNA particle-treated mice.

We next examined macrophage and necrotic core content in aortic root sections. Despite plaque size (Supp. Fig. 4G) and macrophage content (Supp. Fig. 4H-I) being similar between the groups, necrotic cores trended to be larger (p=0.074) and their proportion of plaque area greater in the recipients of the *Fcgr4* siRNA-containing particles (Fig. 4I-K) compared to controls. Furthermore, there was a negative correlation between the abundance of Fcgr4+ macrophages in the plaque or eWAT and plaque necrotic core (Fig. 4L).

These data indicate that eWAT Fcgr4+ macrophages help to reduce plaque necrotic core, likely related to their greater efferocytotic capacity (Fig. 3). More generally these results extend the mechanisms by which eWAT can influence other tissues: in addition to secretion of factors and extra-cellular vesicles (reviewed in (47, 48)), there can also be trafficking of macrophages and other leukocytes.

### Weight regain accelerates atherosclerosis progression and diminishes the content of Fcgr4+ macrophages in plaques and eWAT

Weight loss that is sustained is beneficial in reducing CV risk(5–7), consistent with the effects of stCR in promoting atherosclerosis inflammation resolution (Fig. 1-4). Weight fluctuation, however, is associated with worsening of CVD compared to the maintenance of an obese state (e.g., (11, 49)). This is an important clinical issue because maintaining weight loss is extremely challenging, with >60% of patients regaining weight(50, 51), and many exceeding their starting points.

To investigate the mechanisms underlying increased CVD with weight fluctuation, we extended the stCR studies to include an additional group of weight regain (WR). Similar to our initial study, *Ldlr^-/-^* mice were fed a HFHC diet to establish atherosclerosis and obesity. After 24 weeks, mice were randomized to 4 groups with similar average weights with, once again, the diet composition remaining the same: 1) BL, which was harvested at this time; 2) Progression (PR), which was maintained on *ad libitum* feeding and harvested after an additional 8 weeks; 3) stCR, which was fed 30% fewer calories daily for 2 weeks and then harvested; and, 4) WR, which was put on the stCR regimen identical to group 3 and allowed free access to food thereafter for an additional 6 weeks (Fig. 5A), by which time their weight gain exceeded the BL value (Supp. Fig. 5A). As expected from the data in Supp. Fig. 1, two weeks of stCR reduced body weight by ∼15%, while free access to food following the stCR phase resulted in a 9% increase in body weight compared to the BL (Supp. Fig. 5A). These feeding regimens, however, did not significantly change plasma cholesterol levels (Supp. Fig. 5B). Additionally, while weight loss improved glucose tolerance, WR instead promoted marked glucose intolerance (Supp. Fig. 5C-D) and increased the mass of liver, eWAT, iWAT and BAT compared to BL and stCR mice (Supp. Fig. 5E-H).

**Fig 5:**
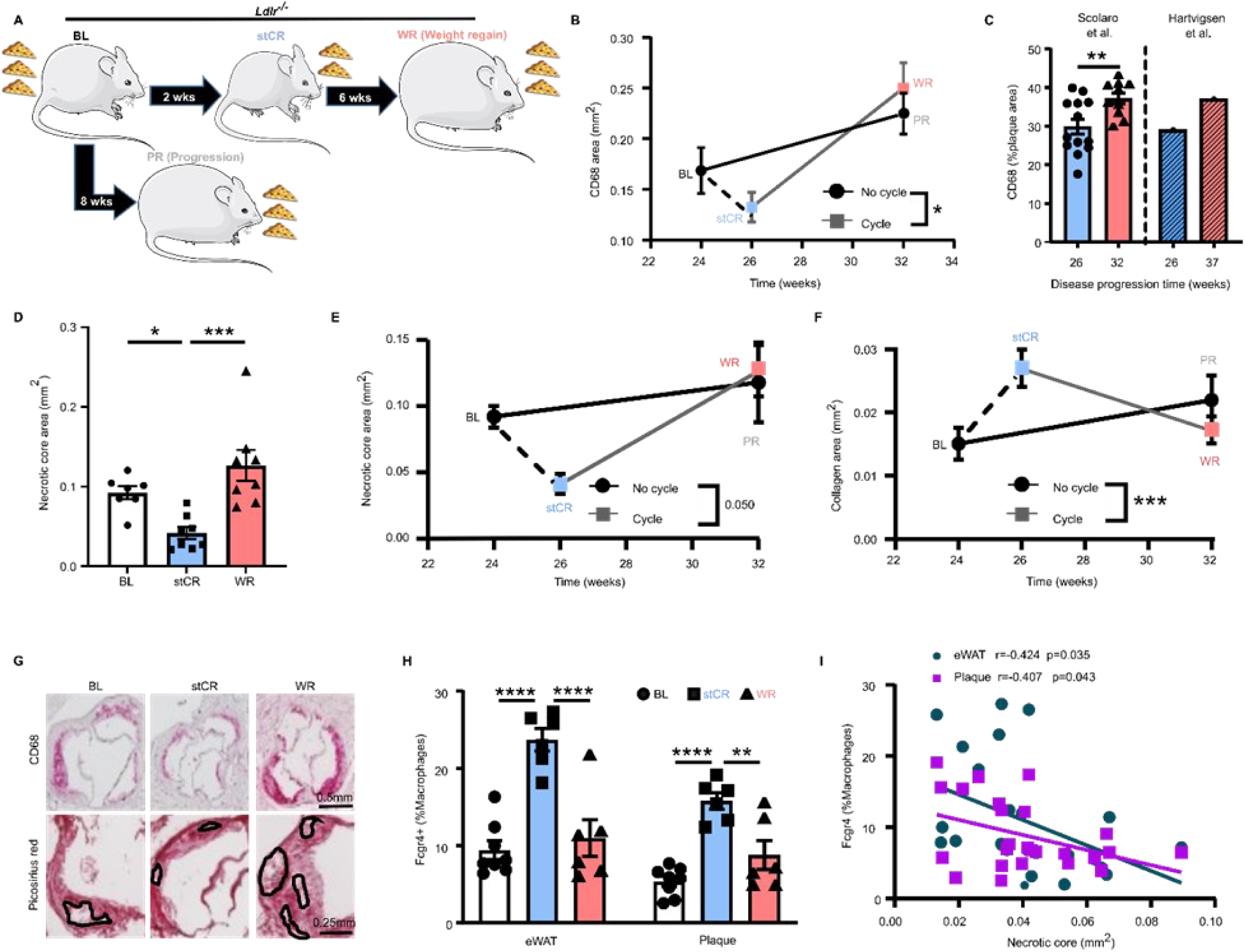
Weight regain accelerates atherosclerosis progression and reverts Fcgr4+ macrophage levels to obese proportions. (A) Schematic of weight regain experiment. Weight regain (WR) was induced by allowing *ad libitum* access to HFHC diet after a two-week weight loss period, achieved by CR. (B) Rate of plaque progression was assessed by comparing increase in macrophage area over time (difference between slopes) between mice that went through weight cycling or that had stable obesity, n=11-15. (C) Side by side comparison between change in plaque macrophage content over time in mice that went through weight cycling and previously published data of plaque kinetics. (D) Necrotic core quantification and (E) rate of change in aortic root sections, n=7-8. (F) Changes in plaque collagen content over time assessed by Picrosirius red positivity in aortic roots (n=7-8), with representative images in (G). (H) Flow cytometry analysis of Fcgr4+ macrophages in eWAT and plaques. (I) Simple linear regression showing correlation between Fcgr4+ macrophages from eWAT and plaques with plaque necrotic core. **\***p<0.05, **p<0.01 ***p<0.001, ****p<0.0001 determined via (B, E, F, I) simple linear regression, (C) two-tailed Student t-test and (D, H) one-way ANOVA with Tukey’s multiple comparisons test.

We next examined the inflammatory state of plaques. Similar to our previous experiments (Fig. 1D-E), stCR reduced plaque macrophage content compared to BL. Notably, WR plaques contained substantially more macrophages than the stCR group (Supp. Fig. 5I). To investigate whether WR alters the rate of atherosclerosis progression, we compared the macrophage accumulation rate in plaques of non-weight cycling mice (i.e., from BL to PR) to those that did have weight fluctuation (from stCR to WR). As expected, macrophages gradually accumulated in plaques of mice fed a HFHC diet without weight fluctuation, resulting in an increase in their area by ∼33% over 8 weeks. Two weeks of stCR reduced plaque macrophage area by ∼28% compared to BL. Strikingly, WR augmented the macrophage plaque area by ∼90% compared to the stCR group (Fig. 5B). Simple linear regression analysis of plaque CD68 area showed a substantial difference in the rates of plaque macrophage accumulation between weight cycling and non-cycling, indicating that WR accelerates atherosclerosis progression.

To independently confirm this acceleration, we compared our data to a published reference, in which Witztum and colleagues carefully assessed atherosclerosis progression weekly in *Ldlr^-/-^* mice(30). In our experiment, plaque macrophage content averaged 29.8% in stCR and increased after only *6* weeks of WR to 37.2% (Fig. 5C). In the aforementioned study(30), plaques had 29% macrophage content after 26 weeks of high-cholesterol diet feeding and reached 37% *11* weeks later (Fig. 5C). Thus, WR caused a 1.8x acceleration in the accumulation of plaque macrophages, in agreement with the difference in slopes in Fig. 5B. Moreover, flow-cytometry analysis of aortic arch macrophages revealed that compared to the stCR group, BL and WR plaque macrophages expressed more of the inflammation-associated marker Ly6C and less of the pro-resolving marker CD206 (Supp. Fig. 5J).

In addition to inflammation-prone macrophage accrual, there were effects on the necrotic core. As shown in Fig. 5D, there was a substantial reduction in plaque necrotic core with stCR compared to either BL or WR. Further comparisons between weight cycling and non-cycling showed acceleration of necrotic core formation with WR (Fig. 5E). Plaque collagen content, which in human plaques is thought to reflect stability(38), showed the opposite trends to macrophage and necrotic core content: plaques from the stCR group contained significantly more collagen than in the BL or WR group (Supp. Fig. 5K); notably, the collagen gain with stCR was lost with WR (Fig. 5F). Representative images of plaques and necrotic cores from BL, stCR and WR are presented in Fig. 5G.

Since Fcgr4+ macrophage accumulation in plaques and eWAT upon stCR was associated with beneficial changes in both sites, we hypothesized that these cells would decrease with WR. To test this, single-cell suspensions were obtained from aortic arches and eWAT, and analyzed for macrophage Fcgr4 levels using flow-cytometry. The results show (Fig. 5H), again, that following stCR there were more Fcgr4+ macrophages in both plaque and eWAT; however, Fcgr4+ macrophage abundances in both sites reverted to their obese, BL, proportions post-WR.

To further characterize the relationship between disease severity and the abundance of Fcgr4+ macrophages, we investigated whether there was a correlation between the amount of Fcgr4+ macrophages in eWAT or plaques with the content of plaque macrophages (Supp. Fig. 5L) or necrotic core (Fig. 5I). The data show a significant inverse correlation between the amount Fcgr4+ macrophages in either tissue and plaque necrotic core (Fig. 5I). A similar correlation was seen with plaque macrophage content, although this did not reach statistical significance (Supp. Fig. 5L).

Taken together, the results show that WR accelerates atherosclerosis progression, with the plaques displaying worse inflammation, enhanced necrotic core formation and reduced collagen content, compared to non-weight cycling conditions. Importantly, Fcgr4+ macrophages in either plaques or eWAT inversely correlated with disease severity in WR.

### Reprogramming of hematopoietic progenitors by WR has durable adverse effects on atherosclerosis

The data thus far show that stCR improves, while WR worsens, atherosclerosis. We previously showed that obese eWAT promotes the expansion in bone marrow of immune progenitors with inflammatory potential, resulting in stimulation of myelopoiesis(4). Thus, we hypothesized that weight loss and regain would also influence the production and inflammatory characteristics of immune cells at the level of the bone marrow. To investigate this, the frequencies of bone marrow progenitors and mature circulating leukocytes in the three conditions were determined. Immune progenitors (including hematopoietic stem cells; HSC, Lin-Sca-1+cKit+; LSK, multipotent progenitors; MPP, Lin-cKit+; LK, common myeloid progenitors; CMP, granulocyte-monocyte progenitors; GMP and megakaryocyte-erythrocyte progenitors; MEP) were least abundant in the stCR group, while the numbers were mostly similar between the BL and WR groups in both frequencies (Fig. 6A) and absolute cell counts (Supp. Fig. 6A). This translated in the circulation to lower myeloid (neutrophils, monocytes, eosinophils and basophils), but higher lymphoid cells in the stCR group compared to both BL and WR (Fig. 6B).

**Fig 6:**
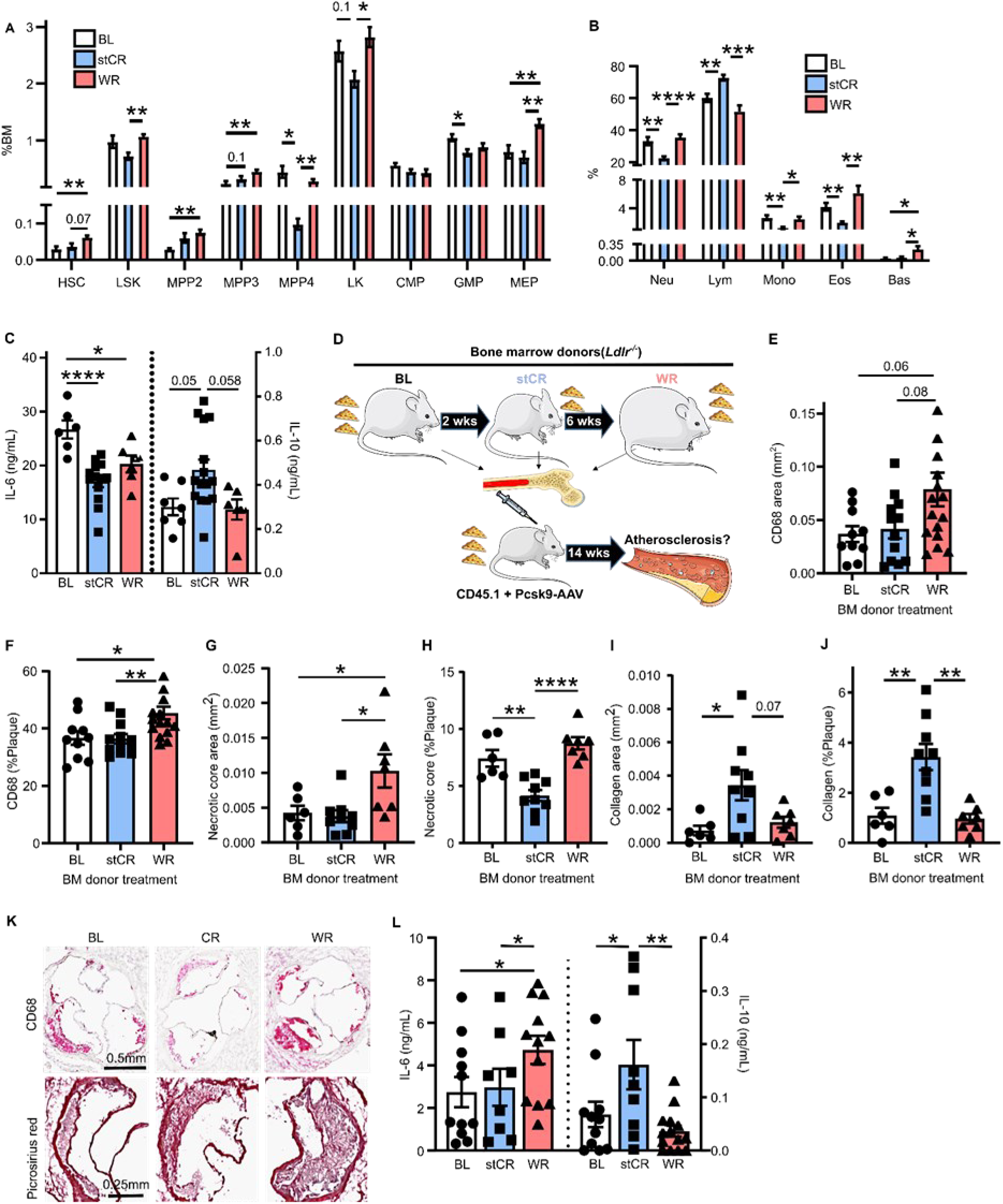
Weight regain induces long-term pro-atherogenic reprogramming of hematopoietic progenitors. (A) Frequencies of bone marrow hematopoietic stem and progenitor cells (n=6-9) and (B) circulating white blood cells (n=13-17). (C) Cytokines produced by bone marrow cells treated *ex vivo* with LPS for 16h. (D) Schematic of bone marrow transplantation experiment. (E) Plaque macrophage content expressed as total area and (F) percent of plaque area assessed by CD68 staining in aortic roots after 14 weeks of HFHC diet. (G, H) Plaque necrotic core and (I, J) collagen quantifications in root sections, with representative images shown in (K). (L) Cytokines secreted by bone marrow cells isolated from bone marrow recipient mice and treated *ex vivo* with LPS for 16h. **\***p<0.05, **p<0.01 ***p<0.001, ****p<0.0001 determined via (A-B) two-way ANOVA and (C-L) one-way ANOVA with Tukey’s multiple comparisons test.

Since there was a substantial decrease in leukocytes with stCR, we investigated whether these immune changes can be driven by eWAT. Hence, mice from the adipose transplant studies mentioned above (Fig. 4A) were analyzed for bone marrow progenitors and mature circulating immune cells. Results show that 2 weeks post-eWAT transplant, there were fewer circulating leukocytes, including neutrophils, monocytes and lymphocytes, in the recipients of the stCR, compared to obese, eWAT (Supp. Fig. 6B). The decrease in circulating leukocytes was accompanied by similar reductions in several hematopoietic progenitor populations in the bone marrow (Supp. Fig. 6C). These data suggest that eWAT following stCR decreases the inflammatory effects of obese adipose tissue on immune progenitors, which would be expected to contribute to beneficial changes in atherosclerotic plaques (Fig. 4C-E).

Because eWAT from mice with obesity or undergoing stCR-induced weight loss influenced circulating immune cells and their bone marrow progenitors (Supp. Fig. 6B-C), we hypothesized that stCR and WR induced innate immune-memory-like changes (aka trained immunity(52)) in myeloid cells and their precursors that contributed to the deleterious effects on atherosclerosis. To test this hypothesis, the responses of bone marrow cells from BL, stCR and WR mice to an inflammatory stimulus (LPS) *ex vivo* were examined. Sixteen-hours post-treatment supernatants were assayed for cytokines classically associated with inflammation (IL-6) and its resolution (IL-10). The results show that cells from the stCR group produced the lowest amount of IL-6 and highest amount of IL-10 compared to cells from BL and WR mice (Fig. 6C). The WR cells also produced less IL-6 in response to LPS compared to cells from BL, but similarly lower amounts of IL-10.

To assess whether these alterations in immune responses are influenced specifically by the eWAT, we performed similar *ex vivo* analyses from bone marrow cells obtained from the fat transplantation study (Fig. 4A). The data show no difference in the production of IL-6, while IL-10 levels were higher in the group that were transplanted with eWAT from stCR mice (Supp. Fig. 6D). This indicates that the stCR eWAT directly affects the inflammatory status of immune cells in the bone marrow by enhancing their production of pro-resolving factors.

Next, we addressed whether these quantitative and qualitative changes to immune cells and their progenitors persist long-term in the context of atherosclerosis and influence plaque properties. To accomplish this, bone marrow transplantation was done, with transfer of cells from the BL, stCR, or WR groups (same mice as in Fig. 5; all CD45.2) to naive, CD45.1 recipients. After recovery from the transplantation, we confirmed substantial bone marrow chimerism with flow-cytometry, using the CD45.1-CD45.2 mismatch (Supp. Fig. 6E). Mice were injected with *Pcsk9*-AAV8 to induce LDLr-deficiency(25), and began HFHC diet feeding to promote atherosclerosis (Fig. 6D).

After 14 weeks, all bone marrow-recipient mice showed similar glucose tolerance, measured by GTT (area under the curve of 29553±2323 in BL, 27995±1728 in stCR and 27362±2175 in WR bone marrow recipients). Plasma cholesterol levels were also similar across all groups (Supp. Fig. 6F). Aortic root tissue sections show that, despite no significant differences in plaque sizes across recipient groups (Supp. Fig. 6G), in recipients of the WR bone marrow, compared to BL and stCR recipients, plaques had more macrophages (Fig. 6E-6F) and larger necrotic cores (Fig. 6G). Although the recipients of BL and stCR bone marrow had similar plaque sizes, macrophage content and necrotic core area, the proportion of necrotic core in plaques was smallest in the stCR recipients (Fig. 6H), which also had the most collagen (as either the absolute area or the % area of the plaque that was positive; Fig. 6I-6J; representative images of plaque macrophages and necrotic cores are presented in Fig. 6K).

Flow-cytometry analysis of aortic arches showed no differences in the proportion of Fcgr4+ macrophages between the 3 groups (Supp. Fig. 6H). This indicates that bone marrow progenitors do not retain long-term memory to produce Fcgr4+ macrophages upon cell transfer. Possibly, transient signals during stCR-induced weight loss promote their appearance, or that if there were epigenetic changes in the precursors, they were not durable, as has been shown in other situations(15, 16). Furthermore, examination of bone marrow immune progenitors (Supp. Fig. 6I-J) and circulating leukocytes (Supp. Fig. 6K) showed no differences in any progenitor or mature cell population across the groups.

Although there were no quantitative differences in the number of circulating immune cells and progenitors, we did find that bone marrow cells from WR recipients had increased IL-6 production in response to LPS *ex vivo* (Fig. 6L). Additionally, the stCR bone marrow retained its ability to produce more IL-10 (Fig. 6L), indicating that hematopoietic progenitors retain some anti-atherogenic capabilities following stCR, mainly influencing plaque collagen content at 14 weeks of HFHC diet feeding. In contrast, there seems to be relatively more pronounced pro-inflammatory reprogramming of immune progenitors following WR, resulting in plaques with enhanced inflammation, larger necrotic cores and less collagen. Taken together, the results suggest that both stCR and WR induce changes in myeloid cells and their precursors that have been seen in other settings (16, 48, 53), particularly the ability to transfer disease-associated phenotypes by bone marrow transplantation.

## DISCUSSION

Obesity increases the risk of atherosclerosis-related CVD(54). Observational studies have accumulated showing that sustained weight loss decreases, while weight cycling worsens CVD risk (e.g., (5, 6, 11)). To isolate the effects of weight cycling on atherosclerosis and to gain insights into underlying mechanisms, we devised a mouse model that allows weight fluctuations without confounding effects from changes in plasma cholesterol or diet composition. From the studies of this model, we obtained the following major findings: 1) stCR induces many features of atherosclerosis resolution, including reductions in the number and inflammatory state of plaque macrophages, increased collagen content, and diminished necrotic cores (Figs. 1 and 5). 2) These improvements are associated with changes in leukocyte sub-population abundances and transcriptomes in both plaque and eWAT (Fig. 2). 3) A distinct macrophage population that accumulates in eWAT following stCR(13) is found in plaques (Fig. 3). These macrophages are distinguished by elevated expression of *Fcgr4* and other phagocytosis-related genes(13). 4) stCR-associated, eWAT-derived Fcgr4+ macrophages in plaques may help clear necrotic cores through their enhanced efferocytosis capabilities (Fig. 4), but upon weight regain, these macrophages disappear from both tissues (Fig. 5). This is associated with accelerated atherosclerosis (Fig. 5), which is consistent with the clinical finding of the aforementioned increased CVD risk with weight cycling. 5) Weight cycling exerts potent effects at the level of bone marrow immune progenitors, as reflected by their altered abundances and long-term reprogramming. This is evidenced in bone marrow transplantation experiments (Fig. 6) in which the plaque phenotype displayed features resembling enhanced stability or disease worsening in recipients of stCR or weight regain bone marrow, respectively (Fig. 6).

Most studies of weight cycling focus on the effects on systemic metabolism (e.g., glucose intolerance and insulin resistance), and include investigations of adipose tissue (55–57) and the liver(58–60). To our knowledge this is the first study looking at concordant changes in eWAT and atherosclerosis with weight fluctuation. Going forward, this mouse model will be a valuable tool to identify further immunological mechanisms that influence atherosclerosis, and possibly other obesity-related co-morbidities in the common situations of weight loss and regain, which is seen in >60% in patients who have dieted(50, 51). Already with a mouse model in which food intake, but not diet composition, is manipulated, we have shown profound short- and long-term changes to the immune compartment and atherosclerosis. The adherence to the same diet composition, with moderate reduction in food intake, is an important distinction of our study compared to others, which involve rounds of diet switching between high fat and normal chow diets (e.g., (12, 16)). Though some diets were shown to promote epigenetic changes and inflammatory reprogramming in immune precursors (e.g., (15, 16)), it is inherently difficult, if not impossible, to discriminate between the effects of the caloric content vs. the components of the diets on the metabolic or immunologic state of the mouse. For instance, Christ et al. showed that Western Diet feeding of *Ldlr^-/-^* mice results in inflammatory reprogramming of immune progenitors, even at four weeks after switching to a chow diet that did not result in significant weight loss (15). Recently, Caslin et al. demonstrated that ATMs derived from HFD-fed mice are more inflammatory than from lean ones, a phenotype that was retained after six weeks of normal chow-induced weight loss(12). Yet another example of the confounding effects of changing diet composition and caloric intake at the same time is illustrated by a recent study in which in a pre-clinical model of macular degeneration, the exacerbation of disease was initially attributed to obesity. In subsequent elegant studies, the causative agent in the diet was identified as stearic acid, suggesting that it was the lower content of this fatty acid in the chow-fed mice, and not their weight loss, that protected them from macular degeneration(16).

In addition to these considerations, there is a fine line separating CR and undernutrition. While the former to a degree is well known to improve health in multiple species, the latter has been shown to promote immune dysfunction (reviewed in (61, 62)). In the first week of diet switch from high fat diet to normal chow, there is a drastic decrease in food consumption, accounting for ∼70% and ∼35% reduction in caloric intake compared to HFD and normal chow-fed mice, respectively(57). This dramatic reduction in caloric intake may induce physiological responses that are similar to starvation or malnutrition, rather than the protective responses attributed to CR. Taken together, the points raised here support the importance in future studies to carefully dissect the differences between various diet regimens to find the most beneficial interventions and their underlying mechanisms.

Our results reveal long-term changes to immune progenitors upon weight loss and regain (Fig. 6). As noted above, recent data indicate that obesity promotes inflammatory reprogramming of macrophages (55) and immune progenitors(16). ATMs from previously obese and weight cycling mice (in a model of diet switching from HFD to normal chow) were shown to retain the hyper-inflamed state in response to *ex vivo* stimulations with toll-loke receptor ligands(55). Our study is in agreement with the weight cycling, but not the weight loss results. We show that both stCR and weight regain bone marrow cells produce less IL-6 in response to LPS than obese-derived cells (Fig. 6C). These inconsistencies between studies might be due to the diet regimens, as indicated above, or the different times used as study endpoints. Nonetheless, our data show that weight regain bone marrow produces elevated IL-6 levels even 18 weeks post bone marrow transplant (Fig. 6L), emphasizing long-term inflammatory effects.

Interestingly, the *in vitro* and *in vivo* data also demonstrate that stCR promotes long term beneficial changes to immune progenitors. *Ex vivo,* bone marrow cells from stCR mice produce more IL-10 (Fig. 6C), even 18 weeks after bone marrow transplant (Fig. 6L). *In vivo*, we see beneficial changes to plaques, mainly increased collagen and reduced necrotic cores (Fig. 6H), in recipients of stCR bone marrows. This suggests that while weight loss may not necessarily overturn the inflammatory programming of bone marrow progenitors from obesity, it might induce pro-resolving features. These results further highlight the importance of retaining weight loss, since weight regain induced the most deleterious changes to immune progenitors (Fig. 6E-K).

As presented in Results, extensive bioinformatic analyses of the leukocyte populations in eWAT and plaques were performed. Of particular interest was the comparison of the transcriptomes of eWAT and plaque leukocytes in obesity and stCR-induced weight loss. Notably, the data show that all immune clusters are shared across these two tissues, albeit in different proportions. It has been long appreciated that tissue microenvironments regulate gene expression of the residing cells (63). This is also reflected in our data, showing mostly non-overlapping gene signatures in eWAT and plaques (Fig. 2C). The data further reflect that transcriptional changes are more coordinated across immune clusters within each tissue compared to cross-tissue changes: we found several genes that coordinately change in multiple (up to 15) clusters within a tissue in response to obesity and stCR (Supp. Fig. 2F and Supp. Table 1). Importantly, as noted above, bioinformatics analysis of this single-cell RNA-seq dataset found the enrichment of a stCR-biased cluster in both plaques and eWAT, namely macrophages distinguished by elevated *Fcgr4* levels (Fig. 2B, 2F).

Our data indicate that the benefits of stCR in atherosclerosis may be attributed, at least in part, to Fcgr4+ macrophages. As just noted, these cells accumulate in both eWAT and plaques upon stCR, but they also disappear after weight regain. An important aspect that remains unknown about Fcgr4+ macrophages is their origin and fate. These macrophages can originate from circulating monocytes that are being recruited to tissues during weight loss. Another possibility is that macrophages that accumulate in tissues during obesity change their function with weight loss and acquire the Fcgr4+ macrophage phenotype. Similarly, their disappearance upon weight regain might be explained by egress of Fcgr4+ macrophages from tissues or their acquisition of a different phenotype in which they lose expression of *Fcgr4*. To answer these and other questions *in vivo*, new genetically engineered mice will be needed.

Our results further suggest that Fcgr4 is not only a marker of macrophages that accumulate with weight loss, but also has functional importance. The expression of *Fcgr4* in eWAT macrophages is associated with reduced plaque necrotic cores, as shown in Fig.4F-L, where local knockdown of *Fcgr4* specifically in eWAT macrophages resulted in larger plaque necrotic cores. The data further suggest that *Fcgr4* expression in macrophages enhances their efferocytotic capability (Fig. 3C-D). It is still unknown, however, how *Fcgr4* regulates efferocytosis beyond its being a receptor that promotes the phagocytosis of immune complexes(64). It is possible, for example, that the expression of *Fcgr4* induces a transcriptional program that facilitates efferocytosis. Another possibility is that *Fcgr4* itself acts as the receptor through which dead cells are being engulfed. If the latter is correct, this will mean that antibody-mediated efferocytosis and macrophage-B cell crosstalk is necessary for this process through antibody production in B cells and engulfment of opsonized particles via Fcgr4 in macrophages. Coating of apoptotic cells by antibodies and their clearance via Fc receptors have been shown in other situations (reviewed in (65)), and we hypothesize this may be a contributing mechanism as well. Future studies will elucidate this issue.

The fact that *Fcgr4* knockdown in eWAT macrophages during stCR impaired the reduction in plaque necrotic core (Fig.4F-L) also suggests an interorgan communication between eWAT and vasculature. It is well established that adipose tissue communicates with many other organs, through secretion of cytokines, adipokines, lipids, metabolites and non-coding RNAs (reviewed in (66)). One mechanism by which adipose tissue communicates with other organs is through the secretion of extracellular vesicles(66). Receptor-ligand expression analysis of plaques and eWAT showed several possible interactions between plaque and eWAT leukocytes (Fig. 2E, Supp. Tables 4, 5). Although this analysis is provocative, it has several caveats. Primarily, we do not have transcriptomic data of non-leukocytes from either plaque or eWAT, and very likely there are intra-tissue interactions with other cell types. Thus, it will be important to dissect experimentally in future studies how weight cycling influences the adipose secretome and how it reaches and affects cells at distant locations. Another mode of inter-organ communication that is suggested by our studies is through cell trafficking. Specifically, the data show leukocyte trafficking between transplanted eWAT and plaques after a transfer of both obese and stCR adipose (Fig. 4A-B, Supp. Fig. 4A). While it is known that macrophages egress from adipose tissue(67), future studies will address whether trafficking from adipose to vasculature is also found in non-surgical protocols by using newly developed methods to fate map cells originating from adipose tissue, and if so, their quantitative contributions.

In summary, we have developed a pre-clinical model of weight cycling that avoids confounding effects of concurrent changes in caloric intake and diet composition, and which recapitulates critical CV features reported in human studies. In this model, there were CV benefits in stCR that were associated with changes in immune progenitors in the bone marrow and mature macrophages in the eWAT and plaques that promoted atherosclerosis resolution. Weight loss also induced the appearance of macrophages in both eWAT and plaque that express *Fcgr4*, which help clear plaque necrotic cores likely through their enhanced efferocytosis capabilities. In contrast, CV benefits observed with stCR were lost with weight regain, with the accelerated atherogenesis attributable to durable inflammatory reprogramming of immune progenitors. Weight regain was also associated with the disappearance of Fcgr4+ macrophages from eWAT and plaques. Together these data suggest that Fcgr4+ macrophages, as well as innate immune memory, are areas in which to base future interventions to promote the benefits of weight loss and prevent the deleterious effects of weight regain on metabolic inflammation, currently major clinical issues in CVD prevention.

## ACKNOWLEDGEMENTS

We thank Jonathan Guzman for his assistance with immunofluorescence staining and imaging. We also thank Mark Graham and Adam Mullick (Ionis Pharmaceuticals) for graciously providing the LDLr ASO. The following Core facilities enabled our studies: NYULMC Flow Cytometry, Genome Technology Center and Histopathology Core. BS acknowledges support from the American Heart Association (863156) and the NIH (HL131481). OT acknowledges support from the American Heart Association (20SFRN35210936) and the NIH (HL131481). MA acknowledges the support of the European Research Council (ERC) under the European Union’s Horizon 2020 research and innovation program (#864788), the Swedish Research Council (#2019-01837) and the Ascending Investigator Grant – Endocrinology & Metabolism (#NNF20OC0060053). AW acknowledges support from the American Heart Association (18POST34080390) and NIH grants HL151963 and HL131481. EAF acknowledges support from NIH grants HL131481 and HL084312, and from the American Heart Association (20SFRN35210936).

## AUTHOR CONTRIBUTIONS

BS, FK, MP, CD, SP, MLG, CAN, ML, OT, CH and AW conducted experiments. Conceptualization, study design and supervision were done by AW and EAF. Data analysis was performed by BS, EJB and AW. MA helped with experimental design and provided key reagents. Manuscript was written by BS, EJB, AW and EAF, with edits from all other authors.

## Notes

Authors declare no conflicts of interest.

### Competing Interest Statement

The authors have declared no competing interest.

