## Supplementary figures for "SHORT-TERM CALORIC RESTRICTION IN MICE PROMOTES RESOLUTION OF ATHEROSCLEROSIS, WHILE WEIGHT REGAIN ACCELERATES ITS PROGRESSION"

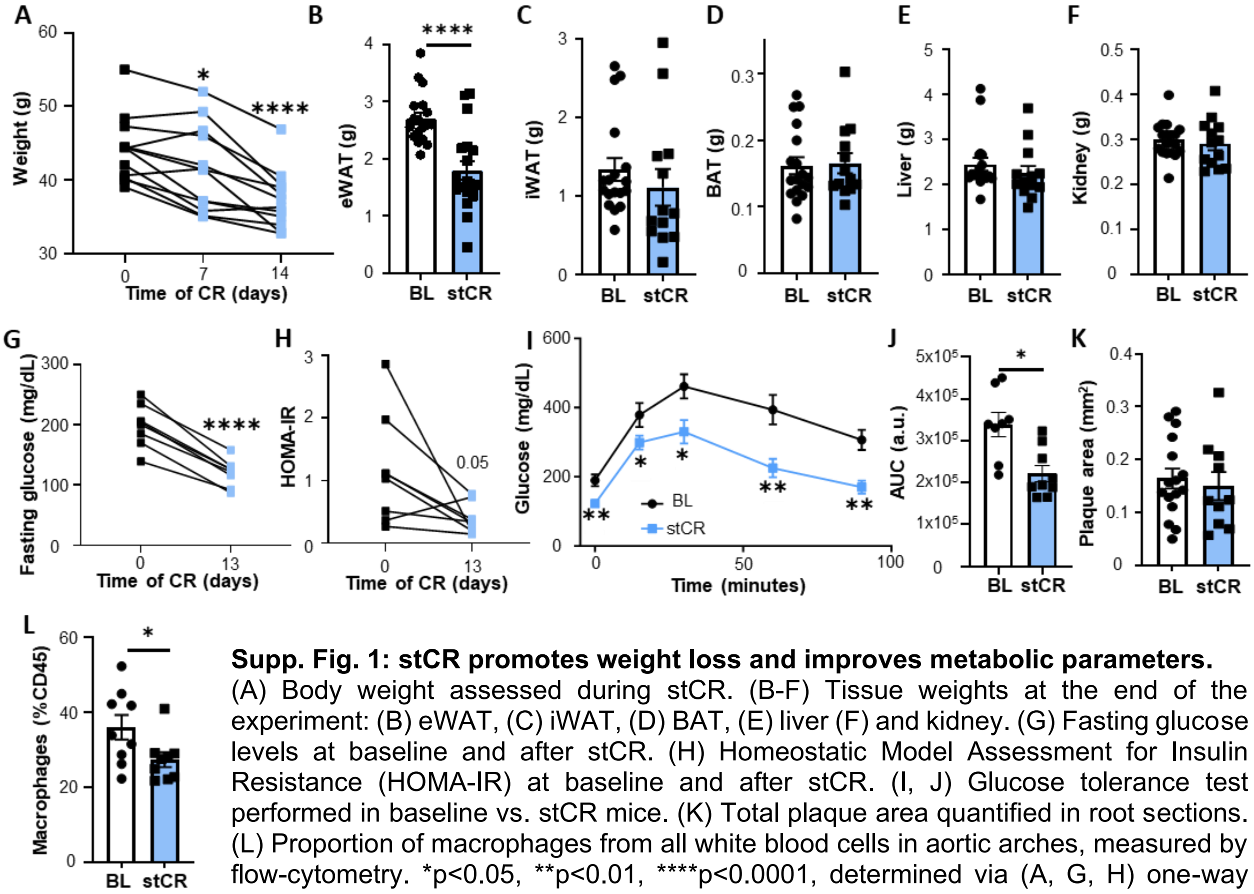


**Supp. Fig. 1: stCR promotes weight loss and improves metabolic parameters.** (A) Body weight assessed during stCR. (B-F) Tissue weights at the end of the experiment: (B) eWAT, (C) iWAT, (D) BAT, (E) liver (F) and kidney. (G) Fasting glucose levels at baseline and after stCR. (H) Homeostatic Model Assessment for Insulin Resistance (HOMA-IR) at baseline and after stCR. (I, J) Glucose tolerance test performed in baseline vs. stCR mice. (K) Total plaque area quantified in root sections. (L) Proportion of macrophages from all white blood cells in aortic arches, measured by flow-cytometry. *p<0.05, **p<0.01, ****p<0.0001, determined via (A, G, H) one-way ANOVA with Tukey’s multiple comparisons test and (B-F, I-L) two-tailed Student's t-test.

**Supplementary Figure 2**

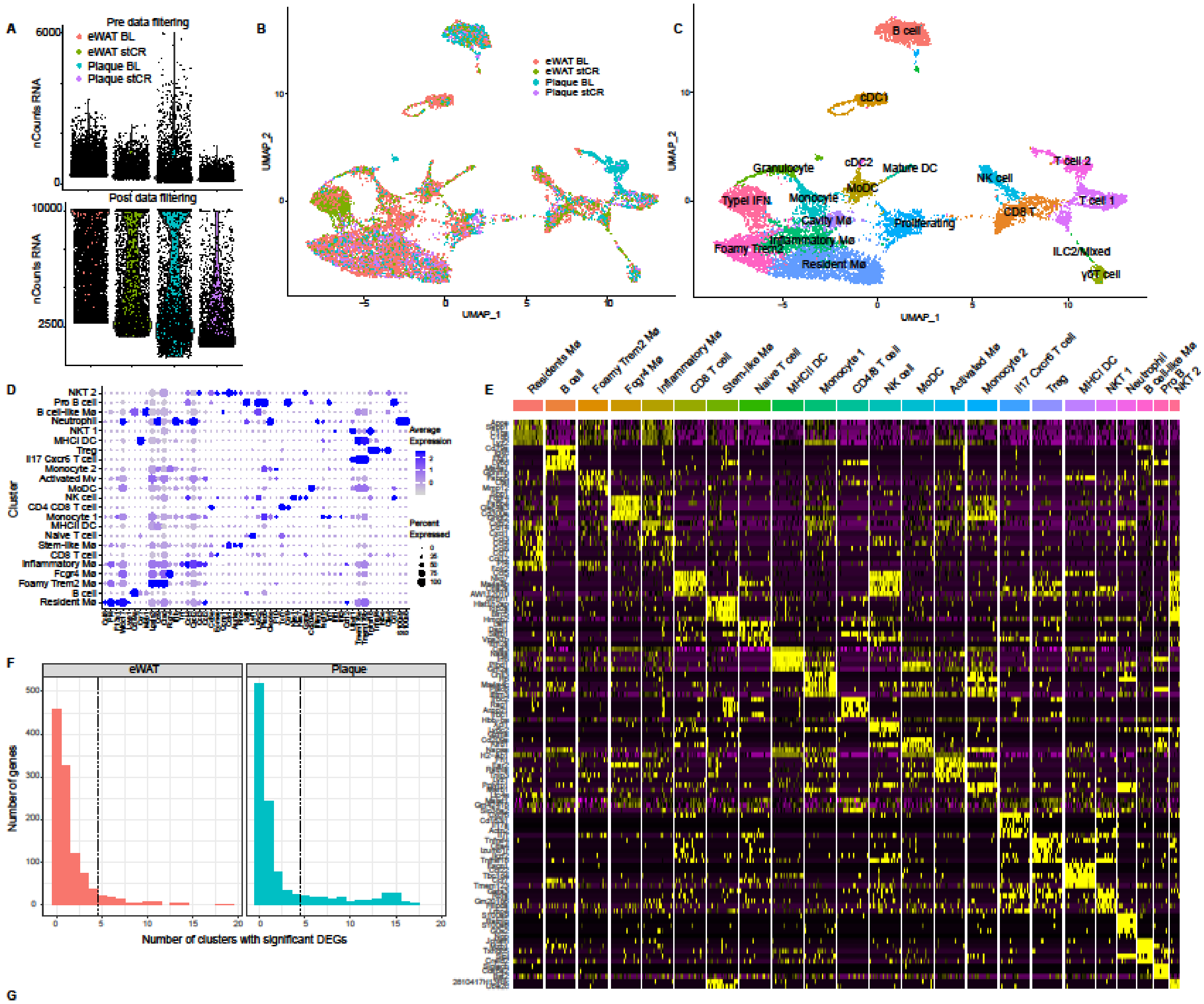


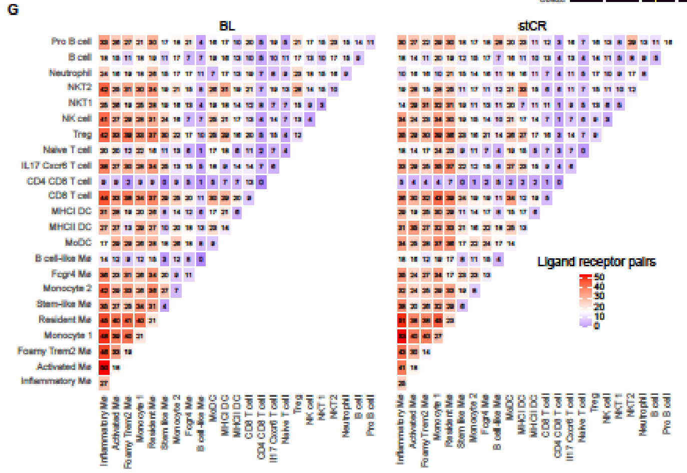


**Supp. Fig. 2: Single-cell RNA-seq of plaque and eWAT leukocytes in obesity and stCR**

(A) Quality control of scRNAseq data pre and post-downsampling for cells containing >10,000 unique RNA reads (see Methods). UMAPs color coded by (B) sample of origin and (C) closest match compared to Zernecke et al.(40) (D) Expression of marker genes defined by Zernecke et al.(40), used to annotate leukocyte clusters in our dataset. (E) Top 5 marker genes per cluster. (F) Number of clusters with shared DEGs (comparing stCR and BL) in eWAT (red) and plaques (blue). (G) Cell-cell interaction analysis, presenting the number of receptor-ligand pairs across all clusters in the eWAT in obese (left) and stCR (right).

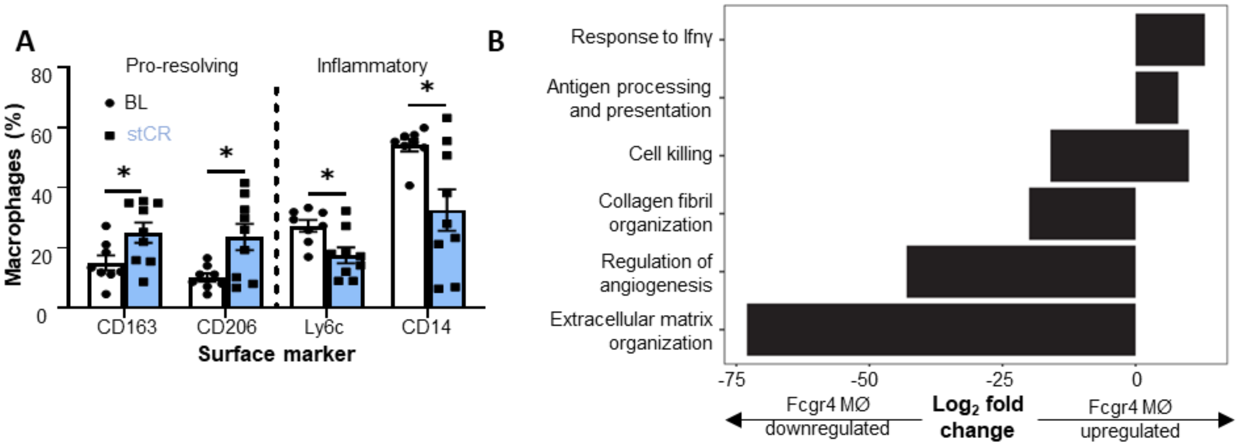


**Supp. Fig. 3: stCR induces a less inflammatory, pro-resolving phenotype of plaque macrophages, partly through induction of Fcgr4+ macrophages**

(A) Flow-cytometry analysis of plaque macrophages after aortic arch digestion. (B) KEGG pathways increased and repressed in Fcgr4+ macrophages from adipose tissue following caloric restriction. *p<0.05, determined via two-tailed Student's t-test.

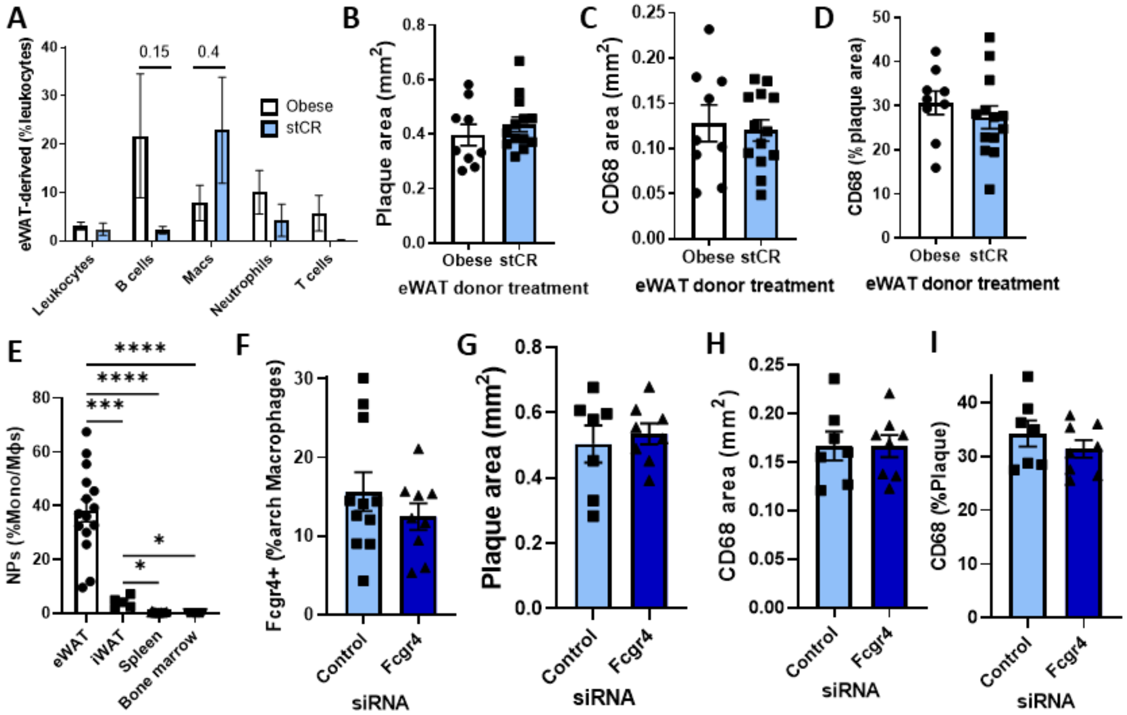


**Supp Fig. 4: Fcgr4+ macrophages from eWAT reduce plaque necrotic core, but do not change plaque macrophage content**

(A) Flow-cytometry analysis of plaque leukocytes of recipient mice, 2 weeks after adipose tissue transplantation, n=5-7. (B) Quantification of total plaque area and (C, D) macrophage content in aortic root sections. (E) Flow-cytometry analysis of particles engulfment by macrophages from different tissues. (F) Flow-cytometry analysis of Fcgr4+ macrophages in aortic arch of mice receiving either control or *Fcgr4* siRNA particles. (G) Quantification of total plaque area and (H, I) macrophage content of aortic root sections, after injections of control or *Fcgr4* siRNA particles. *p<0.05, ***p<0.001, ****p<0.0001 determined via (A) two-way ANOVA with Sidak and (E) one-way ANOVA with Tukey's multiple comparisons test.

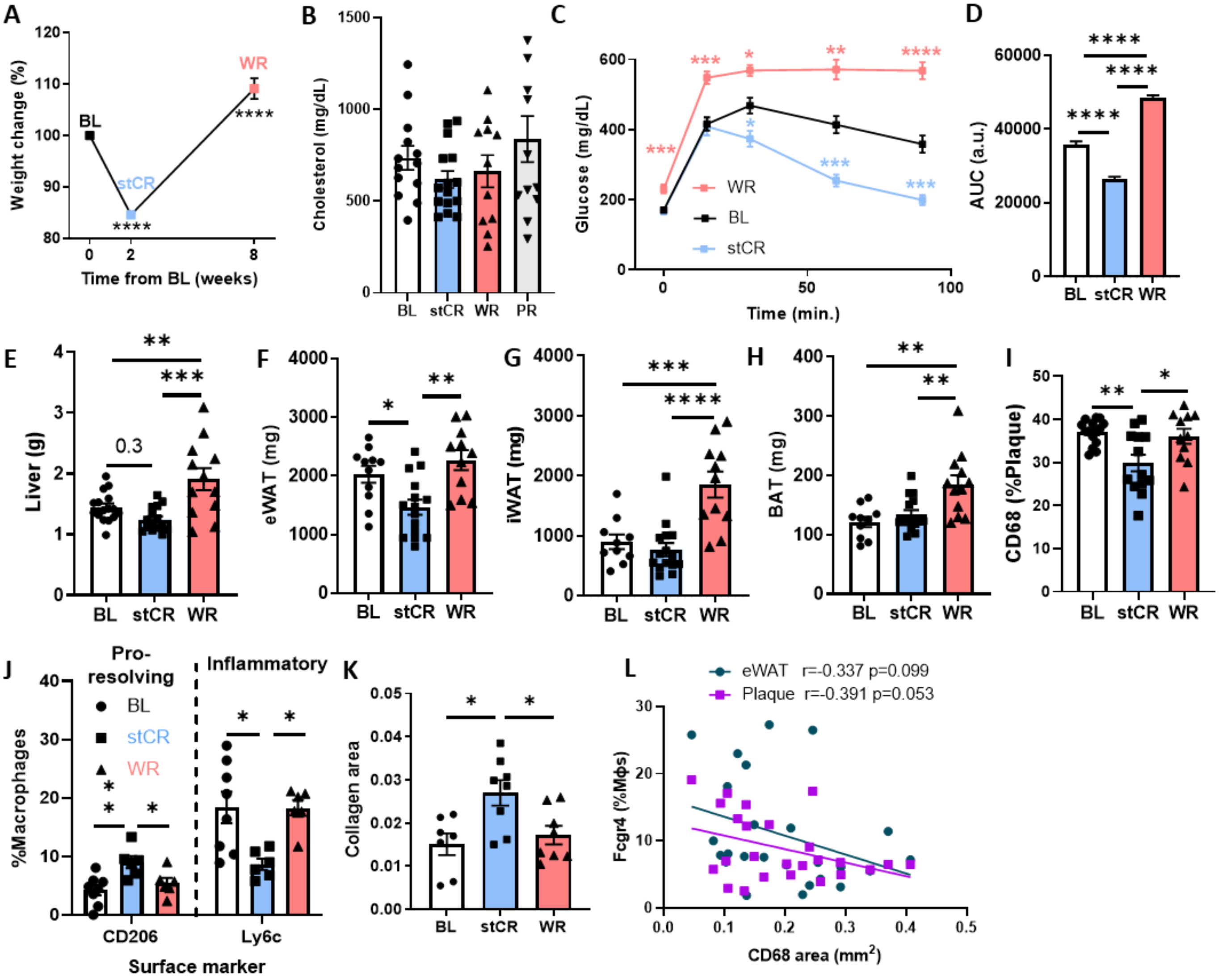


**Supp. Fig. 5: Weight regain worsens metabolic parameters and plaque inflammation**
(A) Percent body weight change during weight cycling, n=29. (B) Plasma cholesterol levels of all groups at harvest. (C, D) GTT performed in baseline, caloric restriction and weight regain groups, n=7-20. (E-H) Tissue weights at the end of the experiment: (E) liver, (F) eWAT, (G) iWAT, (H) BAT. (I) Macrophage percentage of total plaque area measured in aortic root sections. (J) Flow cytometry analysis of aortic arch macrophages. (K) Total collagen content of plaques. (L) Simple linear regression showing correlation between Fcgr4+ macrophages from eWAT and plaques with total macrophage content in plaques. *p<0.05, **p<0.01 ***p<0.001, ****p<0.0001 determined via (A-K) one-way ANOVA with Tukey's multiple comparisons test and (L) simple linear regression analysis.

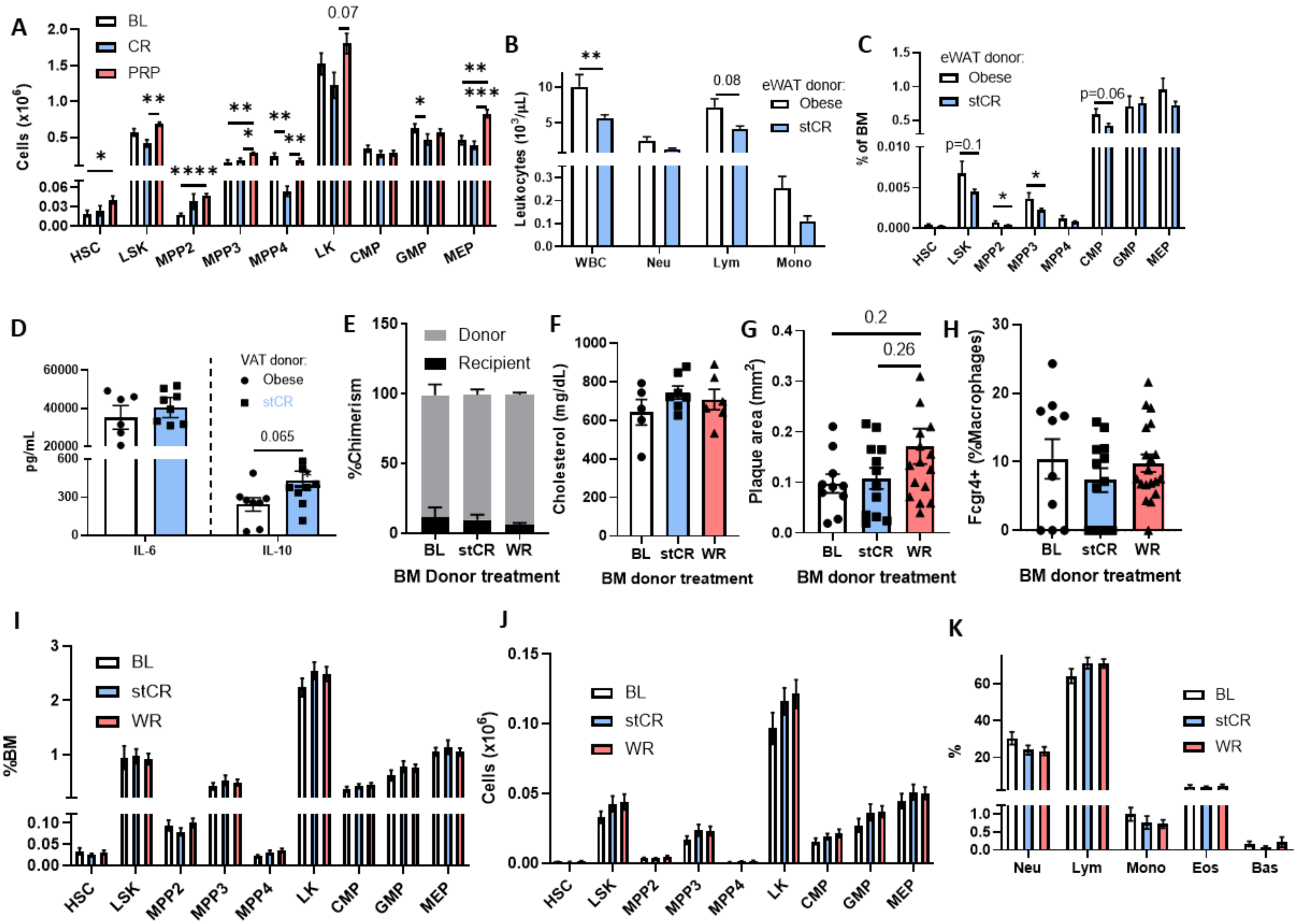


**Supp. Fig. 6: Weight cycling induces long-term reprogramming of hematopoietic progenitors**
(A) Absolute number of bone marrow hematopoietic stem and progenitor cells following weight cycling, n=6-9. (B) Abundances of circulating white blood cells and (C) their bone marrow progenitors in mice transplanted with eWAT from obese and stCR donors (Fig. 4A). (D) Cytokines produced by bone marrow cells harvested from mice transplanted with obese or stCR eWAT and treated *ex vivo* with LPS for 16h. (B-D) n=8-16. (E) Engraftment frequencies of donor cells following BM transplantation, n=6-8. (F) Plasma cholesterol at the end of BM transplantation experiment. (G) Total plaque area of mice transplanted with BL, CR and WR bone marrows and fed HFHC diet for 14 weeks. (H) Flow cytometry analysis of Fcgr4+ macrophages in plaques. (I, J) Bone marrow hematopoietic stem and progenitor cells (K) and circulating white blood cells at the end of the bone marrow transplantation experiment. *p<0.05, **p<0.01 ***p<0.001, ****p<0.0001 determined via (A-C, I-K) two-way ANOVA with Sidak multiple comparison test, (D-E) two-tailed Student's t-test and (F-H) one-way ANOVA with Tukey's multiple comparisons test.
